# Parallel Activation and Interference CRISPR (PAIR) with Sequencing Uncovers DNA Repair Networks Guiding Precision Cell Engineering

**DOI:** 10.64898/2026.05.08.722799

**Authors:** Chen Chang, Dailin Gan, Lu Diao, Lei Wang, Chantale Lacelle, Gary C. Hon, Jun Li, Min Zhao, Siyuan Zhang

## Abstract

The dynamic balance of cellular homeostasis is often maintained by opposing regulatory pathways, yet most genetic screens interrogate them in one direction and therefore miss the bidirectional gene-gene interactions that shape complex phenotypes such as DNA damage response (DDR). Here, we present PAIR (Parallel Activation and Interference CRISPR), a bidirectional perturbation platform that enables simultaneous activation and suppression of distinct genes within the same cell using CRISPR activation (CRISPRa) and Cas13d RNA knockdown. Applying PAIR to the CRISPR/Cas9 induced DSB repair screen, we mapped gene-gene interactions across competing repair branches and identify synergistic perturbations, including NBN activation combined with suppression of end-joining factors, that shift repair outcomes toward homology-directed repair (HDR) and improve the precision of CRISPR-based gene editing. Using coupled PAIR with single-cell transcriptomic, we further demonstrated that NBN activation induces inflammatory and interferon programs, whereas co-suppression of end-joining factors buffers this response, revealing transcriptional states missed by conventional unidirectional perturbations. To translate these findings into non-viral chimeric antigen receptor (CAR) T cell engineering, we developed an mRNA-based strategy for parallel overexpression and knockdown of NBN-anchored DDR effectors in primary T cells, priming the T cells into a transient HDR-favored state that enhances the efficiency of CAR knock-in on the TRAC locus. Together, the PAIR system provides a general framework for studying opposing regulatory networks, uncovering hidden cell states, and guiding cell-state engineering through bidirectional perturbation.

## Introduction

Homeostasis is a hallmark of cell signaling regulation, maintaining stability through the coordinated opposition of activating and inhibitory pathways, this interplay of opposing forces controls cell state stability^1–6^. Thus, mechanistic understanding of the phenotypic consequences of coordinated regulatory pathways is essential for engineering precise cellular state transitions to enhance genome editing outcomes^5, 7^, reprogram cell fate^6, 8–10^, and develop precise therapeutics^11–13^. DNA damage response (DDR) is one of such coordinated regulatory networks where homology-directed repair (HDR) and two competing error-prone pathways, non-homologous end joining (NHEJ) and microhomology-mediated end joining (MMEJ), are each governed by multiple factors^7, 14, 15^. Dissecting these coordinated cells signaling networks requires systematic interrogation of opposing regulatory forces^1, 3, 4, 6, 16^. Although functional genomic studies have broadly dissected DDR gene interactions, most approaches rely on single-gene or dual-gene loss-of-function perturbations within individual pathways and therefore cannot capture the competitive dynamics that arise when HDR and error-prone repair branches simultaneously compete for the same DNA double strand break (DSB)^7, 17–23^. Furthermore, strategies aimed at enhancing precision genome editing must coordinately promote HDR while suppressing competing NHEJ and MMEJ branches to minimize error-prone byproducts – a combinatorial requirement that unidirectional perturbation frameworks are inherently unable to address^20^.

Several approaches have demonstrated bidirectional transcriptional modulation using orthogonal CRISPR systems^9–11, 24–33^. Early strategies employed dual Cas9 orthologs (e.g., *Streptococcus pyogenes* Cas9, *Staphylococcus aureus* Cas9 and Cas12) to achieve independent activation and knock out^31, 32^ or interference^9, 25, 27, 30, 33, 34^. These orthogonal systems have been integrated with single-cell transcriptome profiling: dual-guides CRISPRi^35^, dual-guide CRISPRa^8, 36^ libraries coupled with Perturb-seq mapped pairwise epistasis on transcriptome-level manifolds, parallel activation and interference screens in separate cell populations revealed complementary regulatory circuitry^10, 11, 26^, and most recently, an orthogonal dCas9 system (dSaCas9-VPR and dSpCas9-KRAB) paired with Perturb-seq dissected transcription factor and enhancer regulatory logic within single cells ^25^. Despite these advances, fundamental technical barriers remain. Most platforms, including recent dual-promoter implementations^10, 11, 25, 26^, require multi-promoter architectures to express guides for orthogonal effectors, creating stoichiometric imbalances and complex cloning workflows. Critically, achieving uniform dual perturbations across all cells and incomplete guide capture even in state-of-the-art systems remains significant technical challenges^25^.

Here, we developed PAIR (Parallel Activation and Interference CRISPR), a bidirectional perturbation system that combines CRISPRa and CRISPR/Cas13d-mediated RNA interference within the same cell through a single programmable tandem RNA array. We first showed that the PAIR system can up-regulate and down-regulate different gene pairs simultaneously. Next, we coupled PAIR with pooled PAIR RNA library screening and next-generation sequencing to establish PAIR-seq and applied it to DNA double-strand break repair to map opposing interactions across competing DDR pathways. PAIR-seq identified bidirectional perturbations, including NBN activation combined with suppression of XRCC6, PRKDC, TP53BP1 and POLQ, that shift repair outcomes toward HDR and improve precision genome editing across multiple loci. Furthermore, we integrated PAIR with single-cell transcriptomic profiling to examine the transcriptional effects of NBN-centered perturbations after CRISPR/Cas9-induced DNA damage. scPAIR-seq showed that NBN activation alone induced inflammatory and interferon-associated programs, whereas combining NBN activation with repression of end-joining factors shifted cells into distinct transcriptomic states and buffered these responses. These findings show that bidirectional perturbation by PAIR simultaneously activating one gene while suppressing another - can resolve coordinated interactions between genes acting across distinct DDR pathways. Finally, we translated the DDR rebalancing principles and bidirectional perturbation logic uncovered by PAIR-seq into non-viral chimeric antigen receptor (CAR) T cell engineering by developing POKER (Parallel Over-expression and Knockdown of Effectors via RNA), a bidirectional mRNA-based platform for parallel overexpression and knockdown of DDR effectors prioritized through PAIR validation in a safe and efficient DNA-free format. We show that transient installation of NBN-centered combinations improves CRISPR/Cas9-based CAR knock-in at the TRAC locus during non-viral CAR-T engineering. Together, our work establish PAIR as a general framework for dissecting opposing regulatory networks and translating bidirectional perturbation logic into precision cell engineering.

## Results

### Development of the PAIR system for bidirectional gene regulation

To enable simultaneous gene upregulation and downregulation in the same cell across different gene pairs, we developed the PAIR (Parallel Activation and Interference with CRISPR) system by combing the CRISPR activation (CRISPRa) for upregulation of one gene^37, 38^ and the CRISPR/Cas13d for down-regulation of another gene^38–40^ **(Fig. 1a)**. A single U6 promoter drives a synthetic tandem PAIR RNA (P-RNA) that contains two distinct guide RNA, a Cas9 sgRNA2.0 and a Cas13d pre-crRNA **(Extended Data Fig. 1a)**, the RNA secondary structure prediction suggested that, when arranged in tandem within the P-RNA, the sgRNA and pre-crRNA largely retained their conventional individual secondary structure features^41, 42^ **(Extended Data Fig. 1b)**, which are important for their respective Cas effector functions^40, 43, 44^. Cas13d contains two distinct RNase domains responsible for pre-crRNA processing and target RNA cleavage respectively^40, 45, 46^, enabling it to recognize and process the pre-crRNA array into multiple individual crRNAs for multiplex RNA targeting^47^. We therefore hypothesized that by co-expressing of Cas13, P-RNA and dCas9-vp64/MPH fusion proteins Cas13d would enable a bidirectional regulatory system at same time: Cas13d would recognize and process the pre-crRNA portion of P-RNA to assemble a Cas13d/crRNA RNP for targeted gene knockdown **(Extended Data Fig. 1c)**, while simultaneously releasing a mature sgRNA2.0 that directs dCas9-vp64/MPH to promoter-proximal sites for activation of another gene **(Fig. 1b)**.

**Fig. 1:**
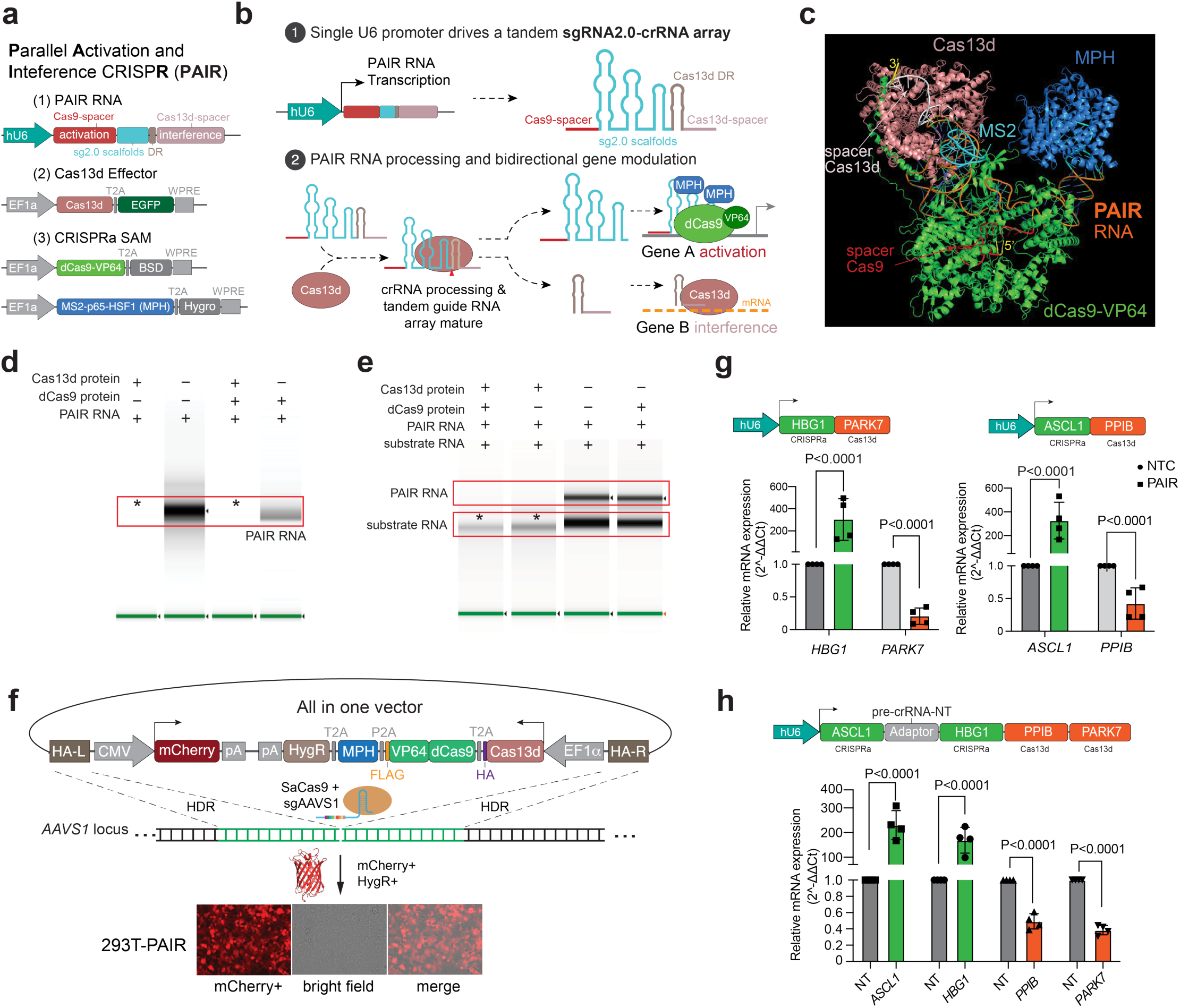
Development of a single-promoter driven PAIR system for coordinated activation and interference of dual genes in single cells. a, Schematics of the PAIR design in which a single U6 promoter drives a tandem PAIR RNA array. **b,** Schematic of expected PAIR RNA processing, enabling parallel gene activation and interference in single cells. **c,** AlphaFold3-predicted structural model of the PAIR RNP complex. The model shows the tandem PAIR RNA positioned between the CRISPRa and Cas13d effector modules, with the 5′ Cas9 spacer and sgRNA2.0 scaffold associated with dCas9–VP64 (green), the MS2 stem loops engaged by the MPH fusion protein (blue), and the 3′ Cas13d spacer/pre-crRNA region associated with Cas13d (pink). PAIR RNA is shown in multicolor, with the annotated spacer and scaffold regions indicating the predicted organization that supports simultaneous CRISPRa recruitment and Cas13d-guided RNA interference. **d,** Cas13d protein processed PAIR RNA during co-incubation with Cas9 protein. The results were replicated in three independent experiments with similar outcomes. **e,** Cas13d protein-mediated cleavage of substrate RNA during co-incubation with PAIR RNA and dCas9 protein. The results were replicated in three independent experiments with similar outcomes. **f,** Schematic of the all-in-one PAIR knock-in system integrated into the AAVS1 locus of 293T cells. mCherry positive cells were isolated by flow sorting following hygromycin selection. **g,** RT–qPCR quantification of simultaneous 2-plex bidirectional gene expression regulation (activation and interference) induced by PAIR RNA constructs in 293T-PAIR cells. **h.** RT-qPCR quantification of 4-plex modulation (two activations and two interferences) by PAIR RNA construct in 293T-PAIR cells. ‘Adaptor’ in schematic indicate pre-crRNA-NT, designed as processing adapter for first two tandem sgRNA2.0. Dots represent biological replicates, and bars represent the mean of n = 4 biological replicates, each with two technical replicates, P values were calculated by one-way ANOVA.

To assess whether the predicted PAIR RNP architecture supports the proposed binding–processing–effector mechanism, we modeled the complex using AlphaFold3^48^. The models indicate that Cas13d and dCas9-VP64/MPH each recognize P-RNA independently and can associate with it simultaneously without steric clash **(Fig. 1c, Extended Data Fig. 1d, e)**. To establish proof-of-concept of the AlphaFold3 predictions, we developed an in vitro transcription (IVT) and RNP assembly assay in which IVT-produced P-RNA (**Extended Data Fig. 1i)** was co-incubated with Cas13d in the presence or absence of dCas9. Following with 30minutes co-incubation, native RNA gel electrophoresis and Tapestation analysis confirmed that Cas13d processed P-RNA equivalently under both conditions **(Fig. 1d; Extended Data Fig. 1f, h)**. Next, to validate whether dCas9 co-incubation affects Cas13d-mediated substrate RNA cleavage, we produced substrate RNA was produced by IVT and co-incubated with Cas13d in the presence or absence of dCas9 for 30 minutes at 37°C. On-target P-RNA reactions produced robust cleavage, whereas NTC P-RNA showed no detectable activity regardless of dCas9 (**Fig. 1e; Extended Data Fig. 1f, g)**. Together, these results confirm that tandem design of P-RNA supports Cas13d-mediated processing and RNA interference activity in the presence of Cas9 protein.

### PAIR enables simultaneous gene activation and interference in cells

Next, to validate PAIR-mediated bidirectional gene regulation in cells under a consistent effector background, we constructed an all-in-one cassette encoding Cas13d, dCas9-VP64, and MPH under EF1α-driven polycistronic expression, flanked by AAVS1 homology arms for site-specific integration (**Fig. 1f)**. Co-delivery with SaCas9 mRNA and an AAVS1-targeting sgRNA^49^, followed by hygromycin selection and mCherry positive enrichment, generated the 293T-PAIR cell line stably expressing all effector components without P-RNA **(Extended Data Fig. 2a, b)**, which was used for subsequent site-specific validation. The conventional dual-guide perturbation for bidirectional^25, 50^ or unidirectional^35, 36, 50^ integrate two orthogonal Cas9 protein and rely on two independent type-III promoters driving orthogonal guide RNAs, requiring multi-step cloning workflows that increase construct complexity, time, and cost **(Extended Data Fig. 2e)**^25, 35, 36^. To simplify the cloning procedure, especially for making multiple cloning more labor and cost efficient, we developed a one-step anneal-and-ligation cloning (OAC) approach to construct the single U6 promoter driven P-RNA expression vectors **(Extended Data Fig. 2c)**. We used the customizable gRNA expression vector (P-CUSTOM)^51^ as the backbone, all P-RNA elements were segmented into multiple complementary single-stranded oligonucleotides (ssODNs) for synthesis, then annealed and ligated into linearized P-CUSTOM to clone individual P-RNA expression vectors. Since P-RNAs share same scaffold^37, 39^, generating a new P-RNA construct only need replacing the 5′ Cas9 spacer and 3′ Cas13d spacer-encoding oligos (**Extended Data Fig. 2d).**

We then selected several gene pairs and constructed P-RNA expression vectors to test whether PAIR system can activate one gene while simultaneously knock down another in the same cells^37, 39^. After 60 hours of expression in 293T-PAIR cells under puromycin selection, we quantified target transcripts by RT-qPCR. Relative to the non-target control (NTC), the PAIR-*HBG1* (CRISPRa)/ *PARK7* (Cas13d) increased *HBG1* expression by more than 300-fold while reducing *PARK7* expression by approximately 80% **(Fig. 1g)**. Similarly, PAIR-*ASCL1* (CRISPRa)/ *PPIB* (Cas13d) increased *ASCL1* expression by ∼180 to 500-fold while reducing *PPIB* expression by more than 50% **(Fig. 1g)**, and PAIR-*IL1B* (CRISPRa)/ *KRAS* (Cas13d) increased *IL1B* expression by ∼200 to 390-fold while reducing *KRAS* expression by 60% **(Extended Data Fig. 2f)**. Together, these results show that the single U6-driven PAIR system enables bidirectional regulation of two distinct targets in the same cells.

Furthermore, we tested whether PAIR could be extended beyond two gene pairs. We built a 4-plex P-RNA cassette and it encoded the validated targets from the 2-plex assays: a tandem array includes two CRISPRa sgRNA2.0 and two Cas13d pre-crRNAs, one extra pre-crRNA with a non-target spacer inserted between two tandem sgRNA2.0 to allow Cas13d to process them as two individual sgRNA2.0 **(Fig. 2f)**. By over-expressing in 293T-PAIR cells, qPCR results showed that the 4-plex design can successfully achieve simultaneously two genes activation and two genes repression, the first activation position target increased by 150∼300-fold, and the second position target increased by 100∼220-fold. Knockdown at the last position reached ∼45% reduction. Considering the total 4-plex PAIR RNA length is over 500nt, we reasoned the decreased efficiency reflects the limited transcriptional capacity of the U6 promoter for long transcript^52^, suggesting that guide placement and transcript length can influence 4-plex PAIR performance in longer U6-driven arrays. In summary, the PAIR system enables efficient opposite-direction regulation of gene pairs using a single U6 promoter. The compact guide-array design makes it straightforward to scale for high-throughput functional CRISPR screen.

**Fig. 2:**
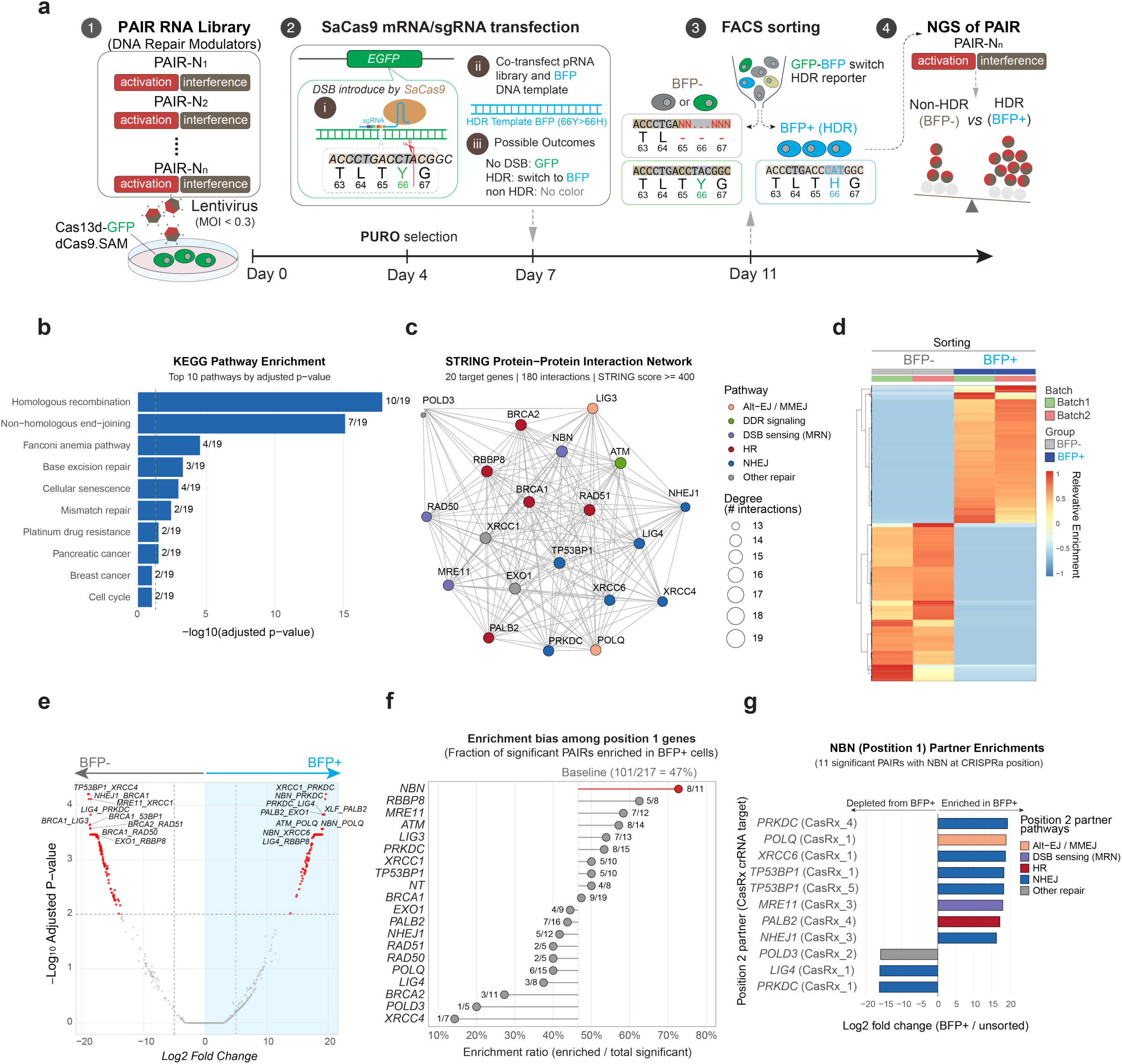
PAIR-seq screen enables high-throughput dissection of competing DNA damage repair (DDR) pathways. **a,** Overview of the PAIR-seq screening strategy. HEK293T reporter cells harboring a GFP-to-BFP HDR reporter were transduced with a dual-modulator PAIR RNA library targeting DNA damage response (DDR) factors. Following SaCas9-induced double-strand break and donor template delivery, cells were sorted based on HDR outcome (BFP+) versus non-HDR (BFP−), and PAIR combinations were quantified by next-generation sequencing. **b,** KEGG pathway enrichment analysis of genes included in the PAIR DDR library, confirming comprehensive representation of major DNA repair pathways, including homologous recombination (HR), non-homologous end joining (NHEJ), Fanconi anemia, and mismatch repair. **c,** STRING protein–protein interaction network of genes represented in the PAIR library, illustrating functional connectivity across DDR modules and supporting the rational design of the combinatorial screen. **d,** Heatmap of significantly differentially enriched PAIR combinations between BFP− and BFP+ populations across biological replicates. **e,** Volcano plot showing log2 fold change versus adjusted P value for PAIR combinations enriched or depleted in HDR-positive cells. **f,** Positional enrichment bias among activation genes (position 1), revealing preferential enrichment of specific HDR-promoting factors in BFP+cells. **g,** Enrichment analysis of position 2 partners paired with NBN (position 1), highlighting coordinated activation–interference interactions across distinct DDR pathways.

### PAIR-seq reveals gene pairs rebalancing DNA repair outcomes

High-throughput CRISPR screens have provided a powerful framework for dissecting DNA damage response (DDR) mechanisms^5, 7, 17, 20, 23, 53^. Most studies have relied on single-gene perturbations^17, 21, 54–58^, and more recent dual-gene approaches have expanded the study of gene–gene interactions (GGI) during DDR^19, 59^. These efforts have also improved precision genome engineering by repressing or inhibiting non-homologous end joining (NHEJ) factors or enhancing homology-directed repair (HDR) effectors^23, 56, 60–73^. However, such unidirectional perturbations does not capture the competition between error-free HDR and error-prone repair pathways such as NHEJ and MMEJ^7, 74, 75^. Consequently, it remains difficult to determine how genes in distinct repair pathways jointly rebalance DNA repair outcomes.^7, 14, 55, 76–81^. To address this gap, we coupled PAIR with synthetic pool library and next-generation sequencing to high-throughput interrogate bidirectional interactions between competing DNA repair pathways^5, 20, 38, 82^.

To apply PAIR as a high-throughput platform for interrogating DSB repair genetic interactions, we generated 20 core DNA repair genes spanning HDR, NHEJ and MMEJ^7, 21, 76, 79, 83, 84^ **(Supplementary Table 4)**. We constructed an unbiased CRISPRa (position 1) × Cas13d (position 2) PAIR library comprising 10,525 bidirectional perturbation combinations, including non-targeting controls (NTC) at each position (**Supplementary Table 5**). Pathway enrichment and protein–protein interaction analyses confirmed that this library broadly captures canonical DDR pathways and their functional connectivity (**Fig. 2b, c**). To functionally score repair bias between HDR and non-HDR at scale, we used Lenti-virus infected GFP positive PAIR effector 293T cells contains harboring a GFP-to-BFP HDR switch^85^, in which a SaCas9/sgGFP-induced DSB at the EGFP (Y66) locus can be repaired using a BFP (H66) HDRT donor **(Extended Data Fig. 3a)**, yielding BFP signal as a direct HDR readout (**Fig. 2a; Extended Data Fig. 3b, c; Supplementary Table 3**). Cells transduced with the PAIR library by MOI<0.3 were transfected with mRNA SaCas9/sgRNA-GFP, and FACS-sorted into BFP+ (HDR+) populations, followed by amplicon sequencing of integrated PAIR cassettes to quantify perturbation combination with each repair outcome (**Fig. 2a; Extended Data Fig. 3 d-f**). Following FACS enrichment and amplicon sequencing, sequencing readouts across biological replicates revealed widespread and reproducible shifts in PAIR representation between BFP⁺ and BFP⁻ populations, and unsupervised clustering of differentially enriched (DE) PAIR combinations robustly separated HDR⁺ from non-HDR samples, highlighting discrete modules consistently enriched or depleted in HDR⁺ cells **(Fig.2 d, e)**.

We next examined whether HDR-associated hits depended on the PAIR positional architecture (CRISPRa target at position 1; Cas13d target at position 2). Significant PAIRs exhibited a pronounced enrichment bias among activation genes (position 1) compared with baseline: 101/217 significant PAIRs enriched in BFP⁺ cells **(Fig. 2f)**. Notably, the top three position-1 activation genes - NBN (NBS1), MRE11 and RBBP8 (CtIP) all converge on DNA end resection^80, 81, 86–88^, a prerequisite early and rate-limiting step for HDR. NBN, MRE11, RAD50 and CtIP to function together to initiate DSB end processing and generate single strand DNA (ssDNA) intermediates. These intermediates are subsequently used by other HDR factors to promote template-dependent repair and can influence competition with end-joining factors at DSB ends^5, 7, 76, 89^. Next, we further annotated HDR-enriched pairs of NBN, MRE11 and RBBP8 by examining their position 2 combinatory partners **(Fig. 2g, Extended Data Fig. 3g, h)**. Among these resection-associated hits, NBN-first-positioned enriched combinations preferentially involved knockdown of genes with established roles in competing end-joining pathways, including canonical NHEJ (PRKDC, XRCC6), MMEJ-associated factors (POLQ) and TP53BP1 consistent with a convergent architecture in which increased HDR capacity is coupled to suppression of non-HDR branches **(Extended Data Fig. 4a)**.

Together, the replicate-consistent enrichment patterns **(Fig. 2d, e)** and positional analyses **(Fig. 2f, g, Extended Data Fig. 3g, h)** nominate bidirectional DDR gene pairs that bias editing outcomes, suggesting a general design principle in which activation of resection-associated HDR factors combined with inhibition of end-joining nodes may shift repair direction toward HDR. Overall, these results establish DDR-PAIR-seq as a scalable framework to map opposing DDR pathway interactions and prioritize candidate combinations for downstream validation in genome editing and cell engineering contexts.

### Bidirectional perturbation of top DDR-PAIR hits enhances HDR-mediated knock-in

To assess the functional consequences of top DDR relevant PAIR-seq hits, we cloned the top-ranked P-RNA into over expression P-CUSTOM vectors^90^ individually and confirmed bidirectional perturbation on the mRNA level by RT–qPCR **(Fig. 3a, Supplementary Table 1)**. CRISPRa-mediated activation of position-1 targets reached 1.6 to 3.4-fold induction relative to the non-targeting control. Cas13d knockdown of position-2 targets reduced transcript levels by 35–75% for most combinations; four pairs showed less than 10% knockdown, likely reflecting local accessibility constraints on Cas13d activity **(Fig. 3b, Extended Data Fig. 4a-c)**. These results indicate that most nominated PAIR combinations produce effective bidirectional perturbation at the RNA level. Next, we tested whether these bidirectional perturbations could shift the repair balance after DSB and improve HDR efficiency^7^. We first co-delivered a mCherry cassette flanked by 1 kb homology arms (HA) with a SaCas9 RNP targeting AAVS1, and quantified integration by junction qPCR across both cassette boundaries **(Fig. 3c)**. Of 21 pairs evaluated, 16 reproducibly increased junction signal relative to the non-targeting control at one or both boundaries **(Fig. 3d)**. The highest-performing combinations shared a common feature: CRISPRa activation of NBN. This pattern held regardless of the paired Cas13d knockdown target, prompting a closer examination of NBN-cantered PAIR configurations. Consistent with DDR PAIR-seq, NBN-centered pairs remained among the strongest performers in our mCherry KI validation. To more precisely quantify HDR versus non-HDR (NHEJ/MMEJ) outcomes after Cas9-based gene editing across different PAIR combinations and multiple loci, we next performed amplicon sequencing at Cas9 cut sites in AAVS1 and CD326 using a 3-bp–barcoded ssODN donor, and quantified HDR: non-HDR ratios using CRISPResso2^49, 91^**(Fig. 3e; Extended Data Fig. 4d-f; Supplementary table 7)**. We first used a dual non-targeting (CRISPRa NTC + Cas13d NTC) as baseline control **(Fig. 3f, g)**, across AAVS1 locus and CD326 locus, Cas13d knockdown of individual NHEJ factors PRKDC or MMEJ factor POLQ generally shifted repair outcomes toward a higher HDR: non-HDR ratio. In parallel, CRISPRa activation of NBN alone also increased the HDR: non-HDR ratio compared with the corresponding NTC conditions. Notably, individually combining NBN activation with four end-joining knockdowns (PRKDC, XRCC6, TP53BP1 or POLQ) produced a stronger enrichment of HDR than perturbation alone, reflected by higher HDR: non-HDR ratios. This enhancement was observed across AAVS1 and CD326 loci. Next, we asked whether simultaneously suppressing all four end-joining factors could further improve HDR or not. To generate a DDR five-factor cocktail, we pooled four individual PAIR constructs, each pairing CRISPRa-NBN activation with Cas13d-mediated knockdown of PRKDC, POLQ, XRCC6 and TP53BP1. This DDR PAIR cocktail configuration achieved HDR: non-HDR ratios above any pairwise combination, demonstrating that PAIR’s crRNA-array architecture can accommodate higher-order combinatorial logic. Together, these results show that systematic DDR-PAIR-seq nominations can be assembled into multi-factor precision-editing strategies, with NBN-centered PAIR designs serving as an effective scaffold for redirecting DSB repair toward HDR.

**Fig. 3:**
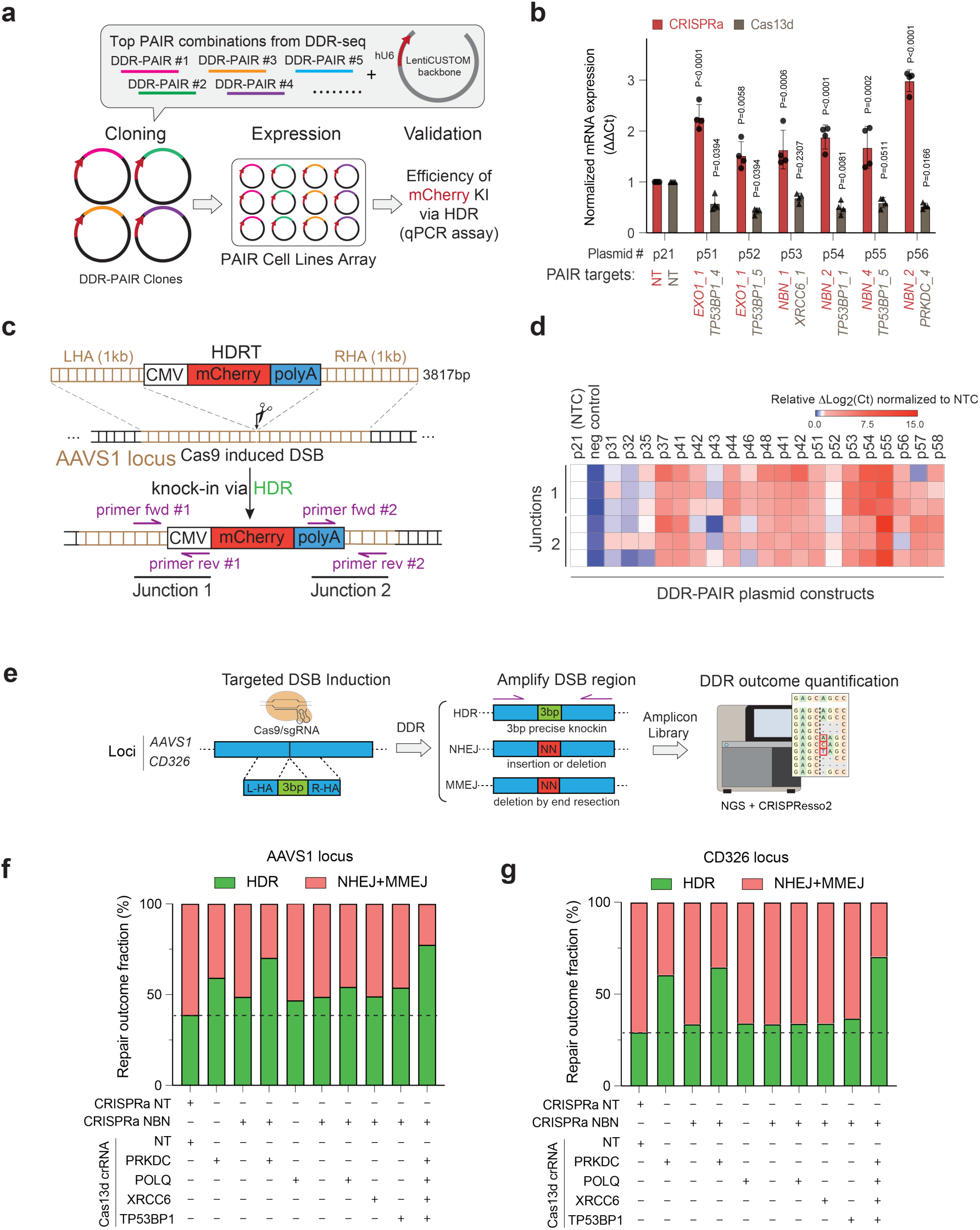
Validation of top PAIR combinations from DDR-seq. a, Schemetics of functional validation of individual top PAIR combinations. **b,** Representative independent validation of DDR-PAIR-seq hits by measuring target gene expressions using RT-qPCR, Dots represent biological replicates, and bars represent the mean of n = 4 biological replicates, each with two technical replicates, p Value is from a one-way ANOVA. **c,** Schematic of CRISPR/Cas9-mediated mCherry HDR knock-in at the AAVS1 locus and quantification of knock-in efficiency by junction genomic DNA qPCR. A Cas9-induced DSB at the AAVS1 locus was repaired using an mCherry HDR template containing 1-kb left and right homology arms. Knock-in was quantified using two junction qPCR assays spanning the 5′ and 3′ integration junctions. **d,** Two independent junction qPCR assays (Junction 1 and Junction 2) were performed separately after genomic DNA extraction to quantify Cas9/AAVS1-mediated mCherry HDRT knock-in efficiency at the AAVS1 locus under the indicated DDR-PAIR conditions (p31–p58), with p21 as the non-targeting control (NTC) and a negative control containing Cas9/sgRNA but no HDRT included. The heatmap shows relative Δlog2(Ct) values normalized to the NTC. n = 3 biological replicates, each measured with two technical replicates. **e,** Schematic of the validation workflow for individual PAIR combinations using NGS-based amplicon sequencing. Following PAIR perturbation and RNP delivery, the targeted DSB region was PCR-amplified and subjected to Illumina sequencing to quantify DNA repair outcomes. Repair categories were classified using CRISPResso2, including homology-directed repair (HDR) and error-prone repair (NHEJ/MMEJ). **f,** Quantification of DNA repair outcome distribution at the AAVS1 locus. Bars represent the percentage of edited reads classified as HDR or error-prone repair (NHEJ/MMEJ). The HDR-to-error-prone repair ratio was calculated from CRISPResso2-classified reads as HDR reads divided by the sum of NHEJ and MMEJ reads. n = 2 biological replicates. g, DDR outcome quantification at the CD326 locus using the same validation assay as in panel f.

### Bidirectional PAIR perturb-seq maps transcriptional manifold in modulating DDR response

Single-cell Perturb-seq bridging defined genetic perturbations to single-cell transcriptomic phenotypes have substantially expanded our understanding of how single gene or dual genes perturbation affect in specific biological processes through the lens of transcriptome manifold^36, 92–94^. To ask whether DDR-PAIR combinations drive an improved HDR through either similar or distinct transcriptional states, we developed single cell PAIR-seq (scPAIR-seq), a bidirectional Perturb-seq workflow that simultaneously captures PAIR perturbation identity (PAIR RNA) and cellular transcriptomes from the same cell. Conventional dual-guide systems based on independent U6 promoters can deliver two gRNAs and have been adapted for single-cell transcriptomic readout, but they remain limited in their ability to directly and consistently recover both guide identities together with mRNA from the same cell^25, 35, 36, 95, 96^. This incomplete linkage between perturbation identity and transcriptional state is particularly restrictive for bidirectional perturbation designs, in which accurate assignment of both perturbation arms is essential. To overcome this, we engineered the 3′ end of the PAIR RNA with a direct capture-compatible sequence (CS1) that is exposed after Cas13d processing and recovered through the 10x Genomics 5′ single cell transcriptome workflow^95^. This design allowed each Gel Bead-in-emulsion (GEM) to recover U6-driven PAIR RNA together with cellular mRNA and hashtag oligonucleotide (HTO) labels, directly reconstructing bidirectional perturbation identity and treatment condition from the same single cell (**Fig. 4a**; **Extended Data Fig. 5a, b**). As an initial proof of concept, we validated the capture strategy in 293T cells expressing four distinct PAIR-CS1 constructs and successfully generated PAIR RNA, mRNA, and HTO libraries, confirming compatibility with standard single-cell library construction (**Fig. 4b, Extended Data Fig. 5c**).

**Fig. 4:**
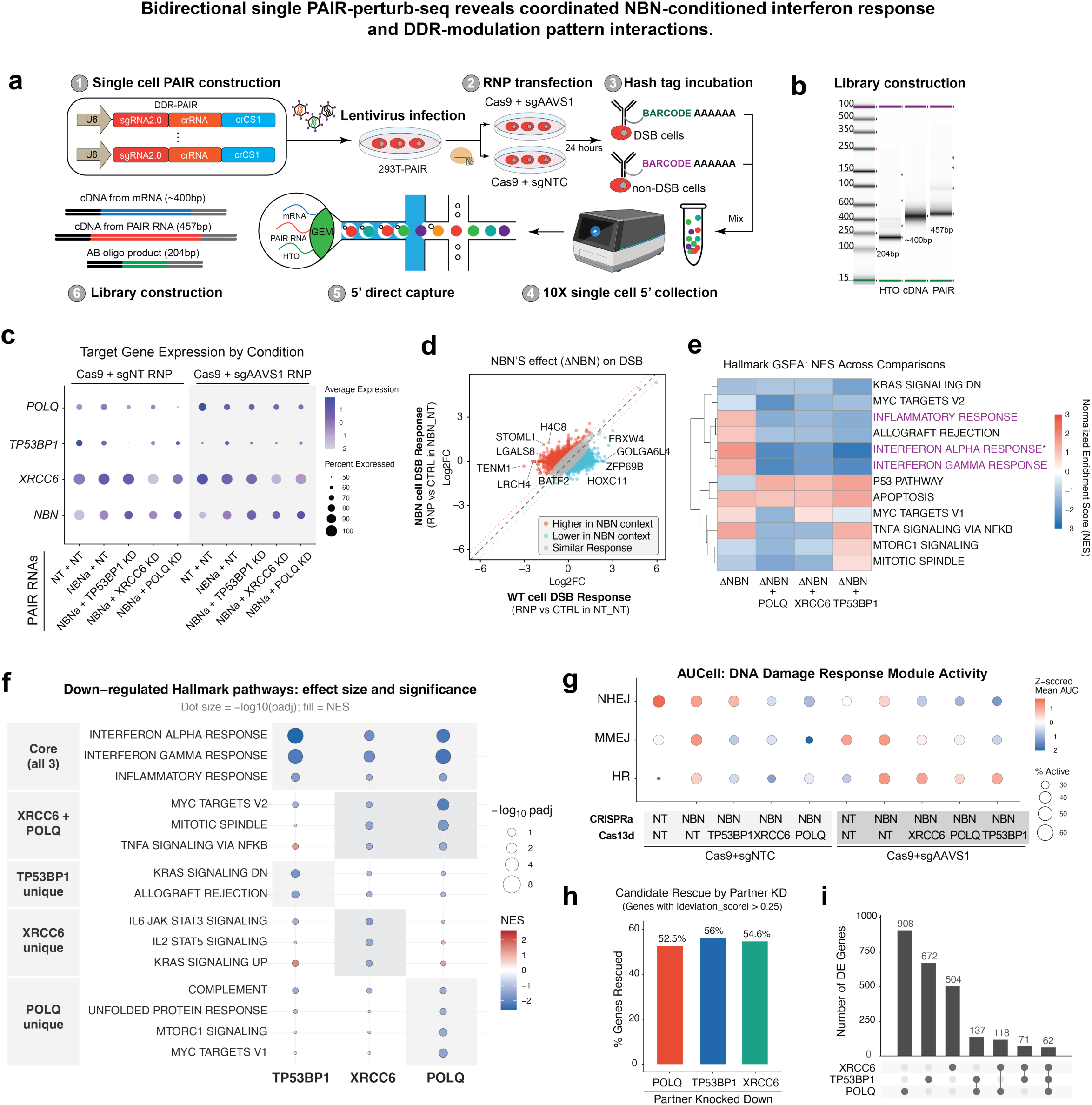
Bidirectional single PAIR-perturb-seq reveals coordinated NBN-conditioned interferon response and DDR-modulation pattern interactions. a, Overview of the scPAIR-seq workflow. 293T-PAIR cells expressing DDR-PAIR constructs were pooled after puromycin selection and divided into Cas9/sgAAVS1 (RNP; DSB-inducing) and Cas9/sgNT (CTRL; no-DSB), hashtag-stained, and profiled on the 10x Genomics GEM-X 5′ platform. Cas13d-processed PAIR RNA with a 3′ capture sequence that is co-recovered with 10X library proprartion pipeline from the same cell. **b,** TapeStation traces of the three pooled single-cell libraries (mRNA, PAIR, HTO) **c,** Dot plot of target-gene expression by perturbation condition across the NT-PAIR and NBN-PAIR arms under both CTRL (Cas9/sgNT) and RNP (Cas9/sgAAVS1) conditions. Dot color, average expression; dot size, percent of cells expressing. **d,** Scatterplot of DEGs in NBN-conditioned DSB response against the wild-type RNP response (log2FC of Cas9/sgAAVS1 vs Cas9/sgNT in the NT_CRISPRa background). **e,** fGSEA normalized enrichment score (NES) heatmap of MSigDB Hallmark pathways across the four pairwise comparisons. Color bar, Noralized Enrichment Score (NES) **f,** Down-regulated Hallmark pathways across the three partner groups, organized as core (shared by all three), XRCC6+POLQ-shared, and partner-unique layers. Dot size, −log10(adj. P); dot color, NES. **g,** AUCell per-cell activity of homologous-recombination (HR), non-homologous-end-joining (NHEJ), and microhomology-mediated-end-joining (MMEJ) gene-set modules across PAIR conditions under CTRL (Cas9/sgNT) and RNP (Cas9/sgAAVS1). Dot color, z-scored mean AUC across conditions; dot size, percent of cells with AUC above the global median. **h,** Barplot showing the fraction of genes recure after partner knockdown down in the NBN-conditioned DSB response set (|score| > 0.25) whose fold change in the dual-perturbation arm is closer to zero than in the NBN-only arm. **i,** Number of differentially expressed genes in each dual-perturbation arm relative to the NBN-only reference (cell-level Wilcoxon rank-sum test, adj. P < 0.05 and |log2FC| > 0.5). Pathway and gene-level statistics are computed on pooled single cells and are used as ranking statistics.

We next applied scPAIR-seq to selected DDR-PAIR hits together with non-targeting controls. Individual PAIR cell lines were pooled after puromycin selection and divided into two treatment groups: one received Cas9/sgAAVS1 RNP to induce a double-strand break (DSB) at the AAVS1 safe-harbor locus, and the other received Cas9/sgNT RNP as a no-DSB control (**Fig. 4a**). Cells were harvested 48 hours after treatment for scPAIR-perturb-seq with and without Cas9-triggered DNA damage. Following quality-control filtering, 2,271 cells were retained across five transcriptional clusters (**Extended Data Fig. 6a, b**). PAIR-mediated bidirectional perturbation was confirmed at single-cell expression level: NBN mRNA was elevated in NBN-CRISPRa-containing groups relative to the NT_NT wild-type baseline, and each partner target transcript was selectively reduced in its corresponding dual-perturbation arm (**Fig. 4c**). At the population level, pseudobulk principal-component analysis across perturbations and treatments confirmed that Cas9-induced DSB remains the dominant axis of transcriptional variation even in the presence of PAIR perturbation, with CTRL and RNP populations separating cleanly along the first principal component across every arm (**Extended Data Fig. 6c**).

### NBN activation potentiates interferon responses under DSB stress

We next asked how NBN activation shapes the transcriptional response. Under the no-DSB condition, NBN activation alone did not produce an appreciable transcriptomic shift across 11,610 tested features (**Extended Data Fig. 6f**). NBN-primed transcriptional identity only emerged once DSB stress was imposed (**Fig. 4d**). To resolve this NBN-primed DSB response, we used a two-step comparison. First, within each genetic background we identified the differentially expressed genes (DEGs) to Cas9/sgAAVS1-induced DSB (RNP vs control) separately in wild-type cells (NT_NT) and in NBN-activated cells (NBN_NT) and identified two independent DSB-response profiles. Second, we directly compared the two profiles to identify genes whose DSB response is shaped by overexpressing of NBN: genes that respond similarly in both backgrounds fall along the diagonal, while off-diagonal genes represent the NBN activation DGE during DSB response (**Fig. 4d**). This analysis reveals that NBN activation selectively amplifies transcriptome response of a subset of DSB-response genes. Within this NBN-primed, DSB-stressed context, we conducted pathway gene set enrichment on the ranked NBN-primed DSB response revealed a diverse elevated pathways that related to interferon, apoptosis, and TNF/NF-ΚB signaling (**Fig. 4e, ΔNBN**). We then asked whether suppression of competing repair partners reshapes the transcriptome manifold. We performed cell-level differential expression of each dual-perturbation arm against the NBN-only reference in the RNP condition and applied pre-ranked fGSEA^97, 98^ against the MSigDB Hallmark collection^99^. All three partner knockdowns produced negative enrichment of broad interferon responses (**Fig. 4e**), suggesting a dominate convergence of suppression of innate-immune signaling. A secondary layer of MYC pathway, Mitotic, and TNF/NF-ΚB signaling dampening was shared between XRCC6 and POLQ but not TP53BP1, suggesting TP53BP1 has functionally distinct roles relative to the other two factors^100^ (**Fig. 4f**).

Knocking down TP53BP1 uniquely attenuated KRAS signaling and Allograft rejection related pathways, while knocking down XRCC6 uniquely affected IL6_JAK_STAT3 and IL2_STAT5 signaling. Knocking down POLQ uniquely dampening MYC, MTORC1 and Complement pathways (**Fig. 4f**).

We applied AUCell^101^, a rank-based enrichment score, using curated homologous-recombination (HR), NHEJ, and MMEJ gene sets^14^ (**Fig. 4g**). HR module activity was elevated in the RNP DSB group relative to its sgNT control group. PAIR NBN pair showed consistent attenuation of NHEJ module activity in the DSB-stress context (negative ΔZ for NBN + TP53BP1, NBN + XRCC6, and NBN + POLQ). Moreover, MMEJ module activity was selectively reduced by POLQ knockdown even at basal damage levels (z =-2.01 in the control), consistent with POLQ’s role as the catalytic core of the MMEJ machinery (**Fig. 4g; Extended Data Fig. 6g)**. The PAIR architecture additionally enables a residual analysis that is unique to bidirectional modulation. Because each cell carries a CRISPRa-activated gene paired with a Cas13d-suppressed partner, we calculated per-gene residual score and epistasis-like interactions^36^. In the RNP-induced DSB context, we found that most of genes fell within ±0.3 of the NBN-only response across the three partner and only a ∼2-4% deviating in both directions (**Extended Data Fig. 6f**), suggesting a non-linear gene–gene interaction effects between NBN activation and suppression of PAIR partner genes.

### No single partner gene knockdown fully reverses the NBN-induced transcriptional state

For each PAIR pairs, we counted the fraction of genes in the NBN-primed DSB response set whose fold change moved toward zero relative to the NBN-only reference after knockdown partner gene - a descriptive measure of candidate rescue. Across the three partners, each single partner knockdown attenuates roughly half (∼50%) of the NBN-primed DEGs after RNP-induced DSB (**Fig. 4h**). Strikingly, despite this convergence on a shared pathway-level endpoint, the three partner-specific gene-hit lists were largely not correlating (**Extended Data Fig. 6e, f**), Even under lenient detection threshold (P < 0.05 and |log₂FC| > 0.3 within DSB-treated NBN-activated cells), the partner-knockdown DE gene sets remained largely non-overlapping: 908, 672, and 504 genes were uniquely affected by POLQ, TP53BP1, and XRCC6 knockdown respectively, compared with only 62 genes shared across all three and 71-137 genes in any pairwise intersection (**Fig. 4i; Extended Data Fig. 6e, h**).

This data indicates that, although several NBN-centered PAIR perturbations similarly favored HDR at the functional level, they were not transcriptionally equivalent at single-cell transcriptome manifold resolution. And each partner coordinately competing DDR pathways reshapes not only IFN inflammatory responses but also the modulate cellular status in respective unique transcriptional state of NBN primed cells, suggesting a transcriptome-level epistasis between genes acting across distinct repair branches.

### POKER enables bidirectional RNA regulation to enhance precise knock-in efficiency

Coordinated NHEJ-HDR balance is not only essential for maintaining genome fidelity, but also play a critical role in CRISPR/Cas-based cell engineering^38, 56, 63, 65, 67, 102–106^. Precise cell engineering is often limited by competing NHEJ and HDR regulators^38, 106^. Our PAIR-seq screening (**Fig. 3**) and scPAIR-seq transcriptome analysis (**Fig. 4)** revealed specific bidirectional modulations of DDR effectors enhance HDR efficiency. This PAIR-mediated bidirectional modulation provided a defined blueprint for optimizing CRISPR/Cas-mediated cell engineering efficiency, where improving HDR while maintaining genome fidelity and high cell viability are highly desirable^5^.

CRISPR/Cas9 based Chimeric Antigen Receptor T (CAR-T) engineering relies on efficient HDR^102, 107–112^. To demonstrate the translational application of the PAIR concept, we explored non-viral CAR-T engineering of primary T cells using a fully programmable and integrated RNA-only PAIR system. We developed POKER (**P**arallel **O**verexpression and **K**nockdown via **E**lectroporation of **R**NA), an mRNA/ crRNA delivery platform that simultaneously overexpresses NBN-Cas13d-GFP hybrid mRNA and an crRNA cocktail targeting four NHEJ/MMEJ regulators (**Fig. 5a**). The POKER system consists of two RNA components produced by in vitro RNA transcription (IVT): a chemically N1-methylpseudouridine (m1Ψ) modified capped Cas13d mRNA that co-expresses the NBN and GFP via a 2A peptides linked overexpression cassette along with a multiplexed pre-crRNA array directing Cas13d-mediated transcript degradation of XRCC6, TP53BP1, PRKDC, and POLQ (**Fig. 5a, b**). Electroporation of the mRNA mixer of modified POKER and GFP produced robust GFP signal in both naïve and activated human primary T cells (**Extended Data Fig. 7a, b**), suggesting that the mRNA is an ideal cargo for gene over-expression in primary T cells. Moreover, capped and m1Ψ-modified POKER yielded substantially higher translation of GFP protein than capped but unmodified controls POKER, despite similar mRNA stability (**Extended Data Fig. 7c, d**). Next, we validated POKER-mediated bidirectional gene regulation in primary human T cells by RT-qPCR. Compared with control Cas13d only mRNA, POKER electroporated primary T cells exhibited robust NBN upregulation alongside efficient and simultaneous knockdown of XRCC6, TP53BP1, PRKDC, and POLQ (**Fig. 5c**). The mRNA-based delivery strategy ensured transient perturbation without permanent genomic modification, and the absence of exogenous DNA avoided the cytotoxic DNA-sensing responses that limit conventional DNA-based DDR modulation approaches^108, 111, 113, 114^ (**Extended Data Fig. 7e, f**). Together, these data show that a single POKER electroporation primes the DDR landscape of primary T cells in the same bidirectional direction identified by PAIR-seq.

**Fig. 5:**
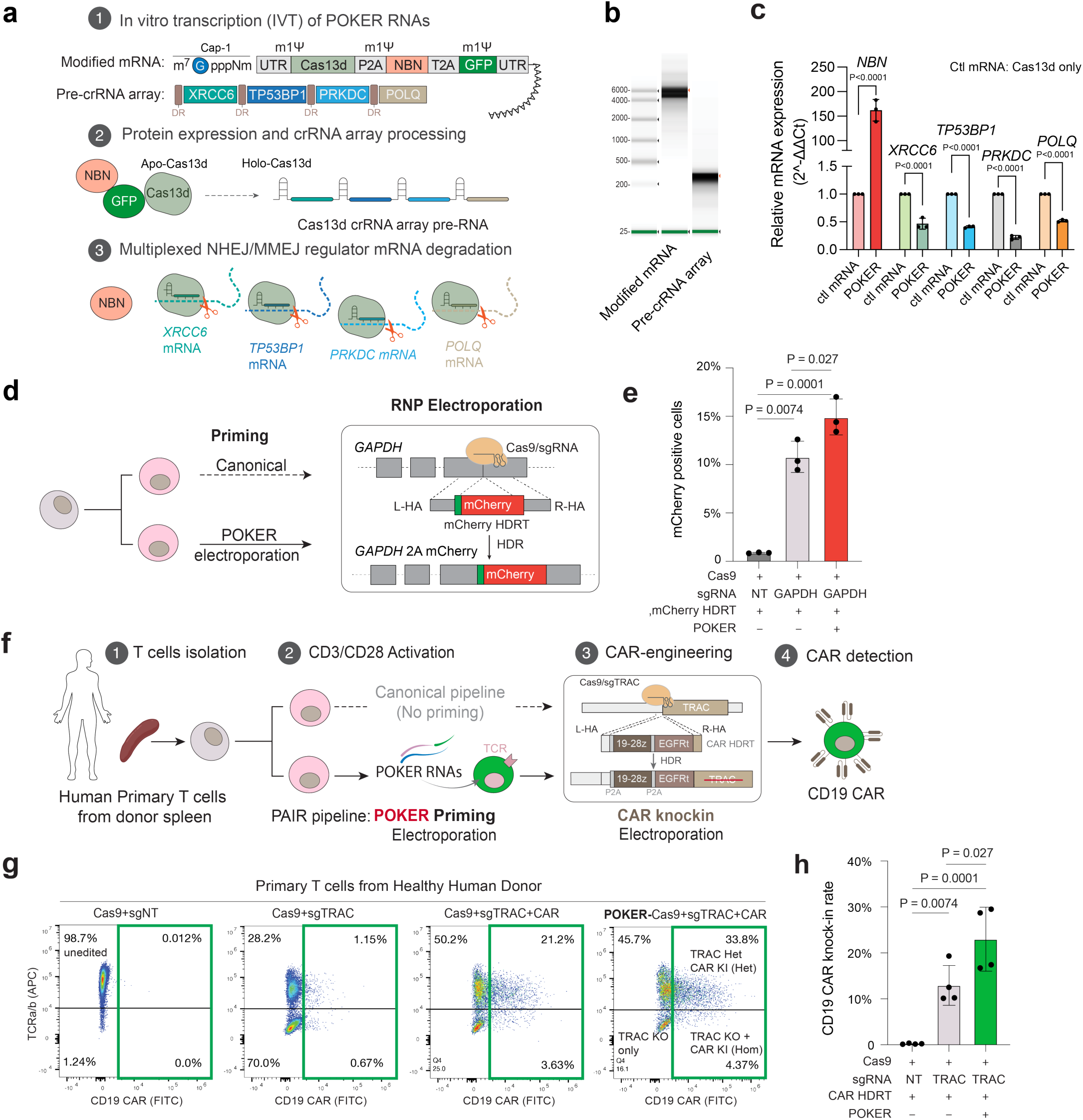
POKER-based bidirectional RNA regulation improves non-viral CAR-T engineering efficiency. a, Schematic of the POKER system. A modified Cas13d mRNA co-expressing NBN via a P2A-linked overexpression cassette is delivered together with a multiplexed pre-crRNA array targeting XRCC6, TP53BP1, PRKDC and POLQ, enabling simultaneous gene overexpression and knockdown within the same cells. **b,** In vitro transcription (IVT) and tapestation electrophoresis analysis of POKER modified mRNA and pre-crRNA array showing expected transcript sizes. **c,** RT–qPCR analysis confirming bidirectional gene regulation in primary human T cells after POKER electroporation, demonstrating NBN upregulation and efficient knockdown of XRCC6, TP53BP1, PRKDC and POLQ. Dots represent independent electroporation replicates from one donor (n = 4), each measured with two technical replicates. Bars indicate mean values. P values were calculated by one-way ANOVA. **d,** Schematic of the GAPDH–2A–mCherry knock-in reporter assay used to quantify HDR efficiency in primary T cells. Cas9/sgRNA targeting the 3′ end of the final GAPDH exon is electroporated together with a dsDNA HDR template encoding an in-frame 2A–mCherry cassette flanked by left and right homology arms. Successful HDR inserts 2A–mCherry at the GAPDH locus, yielding mCherry-positive cells. POKER priming was performed 24 h before Cas9 RNP + HDR template electroporation. **e,** Quantification of mCherry-positive cells in the GAPDH knock-in assay with or without POKER priming. Each dot represents one independent electroporation replicate; bars indicate mean. Statistical significance was assessed as indicated. **f,** Comparison of canonical non-viral CAR-T production and POKER-modified non-viral CAR-T production workflows, illustrating the incorporation of POKER mRNA priming step prior to Cas9/sgRNA-mediated TRAC locus knock-in. **g,** Representative flow cytometry plots showing CD19 CAR knock-in and TCRαβ disruption in human primary T cells following Cas9/sgTRAC editing with or without POKER treatment. Quadrants indicate unedited, knockout (KO), and HDR-mediated CD19 CAR knock-in populations. h, Quantification of CD19 CAR knock-in efficiency across independent electroporation replicates from the same donor under canonical and POKER-modified conditions. Each dot represents one independent electroporation replicate. Bars indicate mean values. Statistical significance was determined by paired two-tailed t-test.

### POKER priming improves the efficiency of non-viral CAR-T manufacturing

To ask whether POKER priming of primary T cell translates into higher HDR-mediated gene knock-in at an endogenous locus, we first established a GAPDH-2A-mCherry HDR reporter assay. A Cas9/sgRNA RNP targeting the 3′ end of the final GAPDH exon was electroporated together with a dsDNA Homology-Directed repair Template (HDRT) encoding an in-frame 2A-mCherry cassette flanked by left and right homology arms (**Fig. 5d**). Successful HDR-mediated integration places the 2A-mCherry sequence in-frame at the C terminus of the endogenous GAPDH coding sequence, enabling detection of precisely in-frame knock-in cells by flow cytometry. POKER was electroporated 24 h before the Cas9 RNP and HDR template delivery as a priming step. Comparing to the canonical knock-in protocol, POKER priming significantly increased the fraction of mCherry-positive T cells around 35% (**Fig. 5e**, p =0.027), indicating that transient POKER-mediated bidirectional DDR reprogramming boosts HDR at a defined endogenous locus in primary human T cells.

We next integrated POKER into a streamlined non-viral CAR-T manufacturing workflow (**Fig. 5f**). In the canonical pipeline, activated human primary T cells are electroporated with a SpCas9/sgRNA RNP targeting the TRAC locus and a dsDNA HDRT encoding a CD19-28z-P2A-EGFRt CAR cassette flanked by TRAC-locus homology arms **(Supplementary table 6)**^111, 115, 116^. We selected the most effective sgTRAC guide from a five-guide screen^111, 115^ (**Supplementary Table 3**) based on TRAC locus cleavage and the efficiency of TCRαβ surface loss in primary human T cells (**Extended Data Fig. 8a, b**). In the POKER-modified pipeline, a single POKER priming electroporation is added one day before the Cas9 RNP + HDRT knock-in electroporation (**Extended Data Fig. 8c**). To evaluate POKER’s effects on CAR knock-in efficiency, we compared the canonical and POKER-modified CAR-T workflows in primary human T cells from a healthy donor by flow cytometry (**Fig. 5g**). In cells receiving the non-targeting control (Cas9 + sgNT), over 98% remained unedited.

Cas9/sgTRAC editing without a donor template achieved efficient TCR disruption (∼70% TCRαβ⁻). In the canonical CAR knock-in condition (Cas9/sgTRAC + CAR HDRT), approximately 21% of T cells achieved CD19 CAR integration, predominantly as heterozygous events (TCRαβ⁻ CAR⁺ Het). POKER pretreatment increased knock-in efficiency. The TCRαβ⁻/CAR⁺ Het fraction increased from ∼21.2% to ∼33.8% (**Fig. 5g**). Quantification across independent replicates confirmed that POKER priming significantly improved CD19 CAR knock-in compared with both the non-targeting control and the canonical sgTRAC-HDRT condition (**Fig. 5h**). Collectively, these results demonstrate that transient POKER-mediated DDR reprogramming translates directly into improved precision CRISPR/Cas9 based knock-in efficacy and CAR integration in clinically relevant primary T cells, establishing a DNA-free priming strategy that enhances non-viral CAR-T manufacturing efficiency without compromising cell viability or manufacturing simplicity.

## Discussion

Here, we establish PAIR, a bidirectional genetics perturbation framework that integrates CRISPRa with CRISPR/Cas13d in the same cell to achieve one the up-regulation and another gene-down regulation. A tandem PAIR-RNA drives both effectors from one U6 promoter, dispensing with the multi-promoter constructs, orthogonal Cas9 platforms, and two-guide cloning steps that have constrained earlier unidirectional and bidirectional designs^25, 36, 93, 94^. Together with PAIR-seq and scPAIR-seq, PAIR is a bidirectional perturbation platform supporting both discovery and engineering applications. PAIR builds on but is distinct from two prior lineages of pooled CRISPR screening. Single-direction pooled screens (CRISPRko, CRISPRi, CRISPRa) and their single-cell genomics platform such as Perturb-seq.

Combinatorial dual-sgRNA libraries and orthogonal Cas9/Cas12 platforms address higher-order genetics but combine perturbations that are uniformly activating or repressing. More recent orthogonal Cas9-based systems, such as CRISPRai^25^, and CRISPR-mediated CDS knock-in combined with shRNA knockdown, as in CRISPRall^24^, introduce bidirectional logic, but at the cost of complex protein engineering or multiple promoter driven orthogonal guide RNAs, which has limited their adoption in large-scale studies requiring cost-efficiency and low labor intensity. The current DDR-based efforts to improve CRISPR gene editing have largely focused on a single direction, either suppressing NHEJ-related genes/proteins^67, 68, 117, 118^ or enhancing/activating HDR-related genes/proteins^56, 65, 66, 104, 119^. However, competing branches of DSB repair (HDR, NHEJ and MMEJ) are not regulated independently, they share substrates, signaling intermediates, and end-state competition for repair-template engagement. Our PAIR-seq data identified that activation of end-resection factors NBN combined with knockdown of end-joining and pathway-choice regulators (PRKDC, XRCC6, POLQ, TP53BP1) improved the higher HDR-to-non-HDR ratio to single direction perturbation of single genes, which broaden the DDR protein relevant CRISPR gene editing improvement.

An important extension of PAIR-seq is its compatibility with single-cell perturbation readouts. By combining PAIR RNA direct capture with single-cell transcriptomic profiling and cell surface hashtag, single cell PAIR-seq enables direct linkage between individual PAIR perturbations and transcriptome state at single-cell resolution, with or without DSB. The single cell PAIR-seq showed that under CRISPR/Cas9-induced DSBs, NBN activation engages interferon-and TNF-driven innate immune signalling, whereas knockdown of XRCC6, POLQ or TP53BP1 buffers these programs. Most interestingly, the genes attenuated by each end-joining partner in combination with NBN are more than 50% non-overlapping, revealing distinct gene–gene interaction logic that would not be captured by single-direction DDR perturbation^21^. Translating these DDR network insights into CRISPR-based primary T cell engineering, we developed an mRNA-based system POKER that utilises PAIR’s bidirectional DDR logic from DDR-PAIR-seq and scPAIR-seq. POKER priming of primary T cells increased non-viral CAR knock-in from ∼21% to ∼34%, demonstrating a potential of PAIR, coupled with POKER, as a general bidirectional perturbation strategy applicable in cell engineering and other biological contexts^10, 11, 28^.

This study has several limitations. First, the PAIR-seq screen used a DDR-related repair-factor library rather than a genome-wide design and therefore provided a rational but not exhaustive map of bidirectional repair interactions in DSB repair. Second, compared with conventional RNA knockdown by CRISPRi, Cas13d offers a compact effector size and the added advantage of guide-array processing. However, knockdown efficiency remains variable across targets. We observed the same pattern during PAIR validation, suggesting that candidates nominated by PAIR-seq require careful follow-up assessment at both the RNA and functional levels. Recent studies further indicate that optimization of the Cas13d effector and delivery conditions may help mitigate Cas13d-associated side effects^120, 121^. Third, although selected combinations were reproduced across orthogonal assays and at more than one locus, the magnitude of benefit will likely remain context dependent across genomic sites, cell types, and donor states. The present scPAIR-seq dataset primarily establishes feasibility and nominates transcriptomic trends associated with bidirectional repair perturbations. Larger-scale studies will be needed to more fully resolve the gene programs, state transitions and interaction structures underlying these effects.

In summary, PAIR provides a scalable framework for interrogating opposing genetic regulations, reading out their combined phenotypic and transcriptomic consequences, and translating selected combinations into cell-engineering strategies. In DNA repair, this approach reveals that precise genome editing can be improved by coordinated modulation of opposing repair branches. More generally, PAIR should enable systematic study of biological processes in which cellular phenotypes emerge from the balance between competing regulation programs.

## Data and Resource Availability

Single cell PAIR-seq datasets with transcriptomic mRNA, PAIR and HTO multiplexing was deposited at GEO: GSE324148 and will be made publicly available upon publishing.

## Declaration of generative AI and AI-assisted technologies in the writing process

During the preparation of this work the author(s) used AI technologies in order to improve writing clarity. After using this tool/service, the author(s) reviewed and edited the content as needed and take(s) full responsibility for the content of the publication.

## Acknowledgments

This study was partially supported in part by NIH grants R01CA255064A1 (to Zhang), 5R35GM145235 (to Hon) and Cancer Prevention and Research Institute of Texas (CPRIT) Scholar Award RR220024 (to Zhang). We additionally acknowledge support from UT Southwestern Simmons Comprehensive Cancer Center. The computational component of this project was supported in part by the BioHPC high-performance computing facility at UT Southwestern.

## Author contributions

Conceptualization and experiment design, C.C. and S.Z.; Data analysis, C.C., D.G., J.L., S.Z.; Experiments and methodology, C.C. with input from D.G., L.D., L.W., C.L., G.H., J.L.

M.Z. and S.Z.; Manuscript writing and revision, C.C. and S.Z.; Study supervision, S.Z. We attest that all coauthors reviewed and pre-approved the manuscript before submission.

## Extended data figure

**Extended Data Fig.1 (related to Fig.1). Establishing PAIR for bidirectional perturbation of two genes in single cells. a,** General sequence of the PAIR RNA. Distinct sequence elements are color-coded, including the spacer, scaffold, and MS2 stem loops in the sgRNA2.0 module, and the direct repeat (DR) and spacer in the pre-crRNA module. The predicted Cas13d processing site within the pre-crRNA region is indicated by a red triangle. **b,** Predicted secondary structures of the full-length PAIR RNA, sgRNA2.0, and pre-crRNA. The PAIR RNA prediction showed preservation of the expected local structural features of both guide modules, including the MS2 stem loops and direct repeat (DR). **c,** c, Schematic of Cas13d-mediated processing of the PAIR RNA. Cas13d (blue) recognizes the pre-crRNA DR region (orange) within the tandem PAIR RNA and processes it at the predicted processing site (red triangle). The processed crRNA then associates with Cas13d to guide target RNA interference, whereas the sgRNA2.0 module remains available for CRISPRa-mediated gene activation. **d,** AlphaFold3-predicted structural model of Cas13d bound to the PAIR RNA. The model shows Cas13d recognizing the 3′ pre-crRNA region of the PAIR RNA, including the Cas13d spacer and direct repeat, while the sgRNA2.0 portion remains positioned outside the Cas13d-bound region. **e,** AlphaFold3-predicted structural model of the CRISPRa module bound to the PAIR RNA. The model shows the 5′ sgRNA2.0 region associated with dCas9–VP64 and the MS2 stem-loop elements engaged by the MPH fusion protein, supporting recruitment of the CRISPRa effector complex through the PAIR RNA. **f,** Agarose gel analysis of in vitro Cas13d-mediated PIAR RNA (P-RNA) processing and substrate RNA digestion. IVT-produced P-RNA was incubated with Cas13d and/or dCas9 protein in the presence of on-target or non-target substrate RNA. Red boxes indicate P-RNA (up) and substrate RNA (bottom) species. **g,** Native PAGE analysis of Cas13d-mediated substrate RNA digesting with on-targeting P-RNA. On-target PAIR RNA supported substrate RNA cleavage in the presence of Cas13d, whereas non-target PAIR RNA or reactions lacking Cas13d did not produce comparable substrate cleavage. **h,** Native PAGE analysis of PAIR RNA processing by Cas13d in the presence or absence of dCas9 protein. Red boxes indicate P-RNA species, showing Cas13d-dependent processing of P-RNA under the indicated RNP assembly conditions. **i,** RNA Tapestation result of T7 IVT generated P-RNAs and substrate RNA.

**Extended Data Fig.2 (related to Fig.1). Establishing PAIR for bidirectional perturbation of two genes in single cells. a,** Flow-cytometry gating strategy used to enrich PAIR-positive HEK293T cells by FACS. **b,** Schematic of the cloning strategy for generating PAIR expression vectors. **c,** Schematics of single U6 promoter driven P-RNA vector workflow. **d,** Schematics of the conventional dual U6 promoter driven dual orthogonal guide RNA vector workflow. **e,** RT–qPCR in 293T-PAIR cells showing Cas13d, dCas9, and MPH expression compared with wild-type HEK293T cells. Points indicate biological replicates; bars indicate mean (n = 4 biological replicates; two technical replicates each). **f,** RT–qPCR showing simultaneous activation and interference by PAIR (IL1B up, KRAS down) in 293T-PAIR cells. Points indicate biological replicates; bars indicate mean (n = 4 biological replicates; two technical replicates each). P values were calculated by one-way ANOVA.

**Extended Data Fig. 3 (related to Fig.2). PAIR-seq screen enables high-throughput dissection of competing DDR pathways. a,** Schematic of the GFP-to-BFP HDR reporter. SaCas9 introduces a DSB at the EGFP(66Y) locus and an HDR donor carrying the BFP(66H) sequence enables template-directed repair, yielding BFP signal. **b,** T7EN1 assay of cutting efficiency, Genomic PCR across the reporter locus following mRNA SaCas9/sgRNA delivery using the indicated sgRNAs (NT, non-targeting control). Red arrowheads indicate the expected amplicon/edited products. **c,** Representative flow-cytometry histograms of BFP signal in cells transfected with non-targeting sgRNA (sgNT) or the indicated sgRNAs together with the HDR donor. **d,** Gating strategy for FACS enrichment of reporter-positive populations. Representative plots and histograms show GFP positive and BFP positive gates used for downstream analysis. **e,** Schematics of DDR-PAIR-seq library structure. **f,** PCR amplification of integrated PAIR-DDR library cassettes from sorted populations (GFP+ and GFP+ BFP+) and control samples, confirming successful recovery of library sequences for NGS. The expected library size is 423bp. **g,** Representative independent validation of DDR-PAIR-seq hits by measuring target gene expressions using qRT-PCR. Dots represent biological replicates; bars indicate mean ± s.d. (n = 4 biological replicates, each with two technical replicates). P values were calculated using one-way ANOVA.

**Extended Data Fig.4 (related to Fig.2, 3). Validation of top PAIR combinations from DDR-seq by amplicon sequencing. a,** Schematic of the two major DNA double-strand break (DSB) repair pathways. Following DSB induction, broken ends are channeled into one of two competing pathways. In non-homologous end joining (NHEJ; top), the Ku70/80 heterodimer rapidly binds DNA ends and recruits DNA-PKcs, followed by the XRCC4–XLF–LIG4 ligation complex, which directly re-ligates the two ends with little or no sequence homology, often introducing small insertions or deletions. In homology-directed repair (HDR; bottom), the MRE11–RAD50–NBN (MRN) complex together with CtIP initiates 5′→3′ end resection to generate 3′ single-stranded DNA (ssDNA) overhangs. The resulting ssDNA is first coated by RPA and subsequently replaced by RAD51, forming a nucleoprotein filament that invades a homologous donor template (HDR template, HDRT) to enable templated, precise repair. **b, c,** Independent validation of DDR-PAIR-seq hits by measuring target gene expressions using RT-qPCR, Dots represent biological replicates, and bars represent the mean of n = 4 biological replicates, each with two technical replicates, p Value is from a one-way ANOVA. **d,** T7 IVT generated sgRNAs for RNP transfection. The expected sgRNA sizes are 96nt. **e,**T7EN1 assay for GAPDH sgRNA cutting efficiency test. **e,**T7EN1 assay for TRAC sgRNA cutting efficiency test. **f,** Representative CRISPResso2 alignment plots showing HDR and error-prone repair outcomes at the AAVS1 locus.

**Extended Data Fig. 5 (related to Fig.4). Design of single cell PAIR-seq library. a,** Schematic of the scPAIR-seq transcript capture and reverse-transcription strategy. The 3′ capture sequence 1 (CS1) was appended downstream of the PAIR RNA to enable direct capture during 10x Genomics 5′ single-cell library construction. After reverse transcription with the feature RT primer and template switching, the final PAIR library contains the cell barcode, UMI, PAIR RNA sequence and capture handle for perturbation assignment. **b,** Sequence of single-cell PAIR RNA. **c,** Schematic of the single-U6-driven scPAIR vector design for direct PAIR RNA capture by 10x Genomics. **d,** Schematic of a conventional dual-U6 promoter strategy for 10x direct capture of dual orthogonal guide RNAs.

**Extended Data Fig. 6 (related to Fig.4). Analyses for scPAIR-seq of the NBN interaction axis. a,** Single cell perturbation assignment, and supporting analyses for scPAIR-seq of the NBN interaction axis. a, Uniform manifold approximation and projection (UMAP) embedding of the 2,271 post-QC cells colored by assigned PAIR identity. **b,** Same UMAP colored by RNP vs CTRL treatment condition. **c,** Pseudobulk principal-component analysis across perturbation arms and treatments; the first principal component separates RNP from CTRL cells across every arm, indicating that Cas9-induced DSB is the dominant axis of transcriptional variation. **d,** Volcano plot of the NBN_CRISPRa vs NT_CRISPRa contrast within the CTRL (no-RNP) condition. No genes pass adj. P < 0.05 and |log₂FC| > 0.5 across 11,610 tested features. **e,** Volcano plots of each partner knockdown (TP53BP1, XRCC6, POLQ) versus the NT reference within the NBN-activated RNP condition. Cell-level Wilcoxon rank-sum; points colored by direction. **f,** RNP-context residual analysis (residual = log₂FC∼AB − log₂FC∼A) per partner. Most genes fall within ±0.3 of the NBN-only fold change (Near-additive residual; TP53BP1 97.6%, XRCC6 98.2%, POLQ 95.8% of 2,000 genes), with a tail of Positive-and Negative-residual genes, enabled by the bidirectional (activation + suppression) geometry of PAIR perturbation. **g,** AUCell mean AUC per condition × treatment (z-scored across conditions) for the HR, NHEJ, and MMEJ gene-set modules. **h,** Jaccard similarity of the partner-knockdown differentially expressed gene sets (cell-level Wilcoxon, adj. P < 0.05, log₂FC| > 0.5).

**Extended Data Fig. 7 frd≥(related to Fig. 5) POKER-based bidirectional RNA regulation improves non-viral CAR-T engineering efficiency. a,** Quantification of mCherry-positive cells in the GAPDH knock-in assay with or without POKER priming. Each dot represents one independent electroporation replicate; bars indicate mean. Statistical significance was assessed as indicated (P values shown). **b,** Representative flow-cytometry gating strategy for analysis of electroporated primary T cells and validation of GFP mRNA delivery, including exclusion of debris and doublets and identification of GFP-positive cells. **c,** Representative fluorescence and bright-field images comparing capped/chemically modified POKER mRNA versus uncapped/unmodified mRNA (with the same pre-crRNA array). Robust GFP signal is observed with the capped/modified POKER mRNA, whereas little to no GFP fluorescence is detected with the uncapped/unmodified construct. **d,** Estimated mRNA half-life in primary T cells for capped/modified versus uncapped/unmodified POKER mRNA constructs (mean ± s.e.m.; P value as indicated), each dots indicate individual donor. **e,** Cell viability of primary T cells following Cas9 RNP electroporation with increasing amounts of dsDNA HDR template (HDRT) across two human donors, illustrating a dose-dependent reduction in viability at higher donor DNA doses. **f,** Representative bright-field images comparing primary T cells after electroporation with dsDNA versus mRNA, illustrating reduced visible cellular stress and improved morphology following mRNA delivery relative to dsDNA under the conditions tested.

**Extended Data Fig. 8 (related to Fig. 5) POKER-based bidirectional RNA regulation improves non-viral CAR-T engineering efficiency. a,** Agarose gel analysis of TRAC locus disruption following electroporation of primary human T cells with SpCas9 RNP and sgRNAs targeting TRAC (representative uncut and indel bands shown), compared with a non-targeting control. **b,** Flow-cytometry screen of candidate sgTRAC guides for efficient TCR disruption in primary human T cells. Representative plots show CD3 and TCRαβ staining following electroporation with the indicated sgRNAs, identifying guides that effectively reduce surface TCR expression for downstream knock-in engineering. **c,** Schematic of the POKER-primed workflow for non-viral TRAC CAR knock-in in primary human T cells, including T-cell isolation and activation, POKER RNA priming, subsequent Cas9/sgTRAC RNP plus dsDNA CAR HDR template electroporation, and downstream CAR detection.

## Methods

### Cell culture and transfection

Cells were maintained at 37°C with 5% CO2 in a humidified incubator and passaged every 2–3 days. Wild type HEK293T cells was cultured in high-glucose Dulbecco’s Modified Eagle Medium (DMEM, ThermoFisher Scientific) supplemented with 10% FBS (Gibico). Cells were split with TrypLE Express (Life Technologies) according to the manufacturer’s instructions. HEK293T cells were seeded on 24-well, poly(d-lysine) plates (Corning) in culture medium. At 80% confluency approximately 12h after plating, cells were transfected with 500 ng of PAIR RNA expression vector in PAIR stable cell-line using 1.5 μl of Calfectin reagent (SignaGen) in DMEM Media (Thermo Fisher Scientific). After 36 hours of transfection, concentration of 10ug/mL of puromycin is added to the transfected samples, 36 hours later the cells are harvested for RNA extraction for qPCR and protein extraction for western blot.

### Construct cloning

For individual PAIR overexpression constructs, the lentiviral backbone pLenti-CUSTOM (addgene#84752), containing a U6 promoter flanked by dual BsmBI restriction sites, was first linearized by BsmBI digestion. The digested backbone was purified by agarose gel extraction using a Zymo DNA Clean & Gel Recovery Kit (Zymo Research). PAIR-encoded single-stranded oligonucleotides (ssODNs) were synthesized by integrated DNA Technologies (IDT). Complementary oligonucleotides were annealed and phosphorylated using T4 Polynucleotide Kinase (New England Biolabs). Insert assembly was performed either by ligation using Solution I (Takara Bio) or by Gibson assembly using the NEBuilder HiFi DNA Assembly Master Mix (New England Biolabs), following the manufacturers’ protocols. Assembled plasmids were transformed into chemically competent E. coli (New England Biolabs) and incubated overnight. Plasmid DNA was isolated using ZymoPURE Miniprep or Midiprep Kits (Zymo Research) according to standard procedures.

### In vitro transcription of PAIR RNA/sgRNA and Cas9 mRNA

For PAIR RNA and sgRNA, a PCR-based approach was used to generate the T7 DNA template. Forward primers containing the T7 promoter sequence were used in PCR amplification. The resulting PCR products were purified using the Monarch DNA Cleanup Kit (New England Biolabs). For T7 in vitro transcription (IVT), 1 µg of T7 DNA template was incubated with the HiScribe™ T7 Quick High Yield RNA Synthesis Kit at 37°C overnight. The IVT products were subsequently purified using the Monarch RNA Cleanup Kit (New England Biolabs). For Cas9 mRNA in vitro transcription, the Cas9 coding sequence (CDS) was first cloned into the mRNA cloning kit (Takara Bio) to serve as the transcription template. The T7 mRNA template was then linearized and purified using a gel purification kit (New England Biolabs) to recover the expected template. 1 µg of the purified T7 template was incubated with the Takara IVTpro™ T7 mRNA Synthesis Kit, supplemented with CleanCap Reagent AG at 37°C overnight. The resulting mRNA was purified using the Monarch RNA Cleanup Kit (New England Biolabs). All IVT RNA products were aliquoted and stored at-80°C for future application.

### 293T-PAIR cell line generation

Generation of the HEK293T PAIR knock-in cell line (293T-PAIR). To establish a stable cellular background for bidirectional regulation, an all-in-one PAIR effector cassette was integrated into the AAVS1 locus in HEK293T cells via CRISPR/Cas9-mediated, homology-directed repair (HDR) (Fig. 1f). The HDR template (HDRT) encoded EF1a-driven Cas13d, dCas9-VP64, and an MPH fusion, separated by three individual 2A peptides, together with a hygromycin resistance cassette and a CMV-driven mCherry reporter encoded on the opposite strand. HEK293T cells were co-transfected with the HDRT and SaCas9/sgAAVS1. Transfected cells were subjected to hygromycin selection followed by mCherry-based enrichment (Supplementary Fig. 1h). Effector expression in the resulting 293PAIR line was quantified by RT–qPCR and compared with wild-type HEK293T cells (Supplementary Fig. 1i).

### Lentivirus production

HEK293T cells were transiently transfected with lentivirus constructs, and packaging plasmids pCMV-dR8.91, and PMD2.G. Lentivirus was collected 48 hours after transfection and filtered through 0.45μm filters. When necessary, virus supernatant was centrifuged at 1500g for 30 min at 4°C to collect virus pellets. The

### In Vitro PAIR RNA Processing and Substrate RNA Digestion Assays

Cas13d and Cas9 proteins were purchased from Molecular Cloning Laboratories Ltd. For PAIR RNA processing assays, reactions were performed at 37°C for 1 hour in RNA processing buffer (20 mM HEPES, pH 6.8; 50 mM KCl; 5 mM MgCl₂; 5% glycerol). The reaction was then stopped by incubating at 75°C for 5 minutes. Samples were analyzed using 8% native PAGE (0.5× TBE buffer), followed by Toluidine Blue O staining (Chem-Impex) or processed using Agilent TapeStation with the High Sensitivity D5000 Kit (Agilent).

### Total RNA extraction and RT-qPCR

Total RNA was extracted using the Quick-RNA Miniprep Kit (Zymo). To eliminate DNA contamination, all samples underwent an additional DNase I (NEB) digestion step. RNA concentration was measured using a NanoDrop spectrophotometer (Thermo Scientific) and assessed by agarose gel electrophoresis with AMD-1000 gel stain (Gel Ltd.). For cDNA synthesis, 1 μg of total RNA was reverse transcribed using the PrimeScript RT Master Mix (Takara). Quantitative real-time PCR (RT-qPCR) was performed using the iTaq Universal reagent (Bio-Rad) on a CFX Opus Real-Time PCR System (Bio-Rad). β-ACTIN and GAPDH were used as reference genes for normalization. Each run included four biological replicates, with two technical replicates per sample. mRNA expression levels were quantified using the ΔΔCT method, and data analysis was conducted using Excel (Microsoft) and Prism 10 (GraphPad). Primer sequences for qPCR are listed in Supplementary Table 2.

### DDR library construction and DDR-PAIR amplification

The DDR library consists of 10,525 PAIR oligos, synthesized by Twist Bioscience. The oligos were cloned into the lenti U6 PAIR backbone using the BsmB1v2 Golden Gate Assembly Kit (NEB). The assembled products were electroporated into competent cells (Biosearch), plated onto 15 cm agar plates containing ampicillin, and incubated at 20°C for 24 hours. Single colonies were harvested in LB medium, and plasmid DNA was extracted using the Plasmid Maxiprep Kit (Zymo) for lentivirus packaging. For DDR Lentivirus Production and Infection, HEK293T PAIR cells (a stable cell line expressing Cas13d and CRISPRa) were cultured in 10 cm dishes. The DDR library (including 10,525 different PAIR RNAs, including negative controls) was packaged into lentivirus and used to infect 293T PAIR cells at a MOI of 0.3. After 48 hours post-infection, 10 µg/mL puromycin was added for selection. The medium was replaced once blank control cells had died. After 7 days of drug selection, DDR-infected cells were split into three groups and transfected with SaCas9/sgRNA and a BFP DNA donor. Flow Cytometry and Sample Collection: at 4 days post-transfection, flow cytometry was performed to sort BFP+ cells. Different cell populations were harvested for library construction: 1.

Sorted BFP+ cells. 2. Unsorted cells. 3. Puromycin-selected cells. Genomic DNA was extracted using the Quick DNA Midiprep Kit (Zymo) for each group. Indexed libraries were prepared for each sample, pooled at an appropriate ratio, and sequenced on a NovaSeq X system (Novogene).

### mRNA vector cloning and in vitro transfection

CDS fragments were synthesized by Twist Bioscience and cloned into the Takara IVT template vector using the Cloning Kit for mRNA Template (Takara Bio, Cat. #6143). CDS fragments were assembled into the pre-linearized vector using In-Fusion Snap Assembly Master Mix included in the kit. Briefly, 50 ng linearized vector, 100 ng CDS fragment and 2 μl 5× In-Fusion Snap Assembly Master Mix were combined in a 10 μl reaction and incubated at 50 °C for 15 min. The reaction was transformed into competent E. coli, plated on LB-kanamycin agar, and plasmids from individual colonies were sequence verified. For IVT, sequence-confirmed plasmids were linearized downstream of the poly(A) sequence, purified, and used as templates for T7 transcription. mRNAs were generated using CleanCap co-transcriptional capping chemistry and complete substitution of UTP with N1-methylpseudouridine-5′-triphosphate. After IVT, mRNAs were DNase-treated, purified, quantified by spectrophotometry, and assessed for transcript size and integrity before electroporation.

### Primary T cell isolation and culture

Primary human T cells were isolated from healthy human donor spleen tissue. Spleen tissue was mechanically dissociated and washed with PBS to generate a single-cell suspension. splenic mononuclear cells were enriched by density-gradient centrifugation using Lymphoprep, and T cells were subsequently isolated using the Pan T Cell Isolation Kit, human (Miltenyi Biotec, Cat. #5250102507), according to the manufacturer’s instructions. Isolated T cells were cultured in X-VIVO 15 medium (Lonza) supplemented with 5% fetal bovine serum (FBS; VWR), 50 μM 2-mercaptoethanol, and 10 mM N-acetyl-L-cysteine (Sigma-Aldrich). One day after isolation, T cells were activated with anti-human CD3/CD28 activator (Stemcell, Cat# 10971) in presence of 25ul/mL, ratio in the presence of 5 ng/ml IL-7 (Biolegend), 5 ng/ml IL-15 (Biolegend), and 300 U/ml IL-2 (Biolegend), exchange medium one time per day. After 2 days of activation, exchange normal culture medium without activator, and T cells were maintained in medium containing 300 U ml⁻¹ IL-2. Medium was replaced every other day, and cells were maintained at 0.5–1 × 10⁶ cells/ml. All T cells were grown at 37 C and 5% CO2.

### T cell electroporation and CAR-T production

HDRT for T cell were synthesized by Twist Bioscience. Primary human T cells were electroporated using the Neon Transfection System (Thermo Fisher Scientific, Cat#MPK5000). sgRNAs were generated by in vitro transcription from PCR-amplified DNA templates, purified, and used for RNP assembly. For each reaction, Cas9 RNPs were assembled by incubating 1 μg IVT sgRNA with 3 μg TrueCut Cas9 Protein v2 (Thermo Fisher Scientific, Cat# A36496) and 1ug HDRT at 37 °C for 15 min, activated T cells were washed once with PBS and resuspended in Neon Resuspension Buffer T with Cas9 RNP/ HDRT immediately before electroporation. For mRNA electroporation, activated T cells were mixed with 2 μg IVT mRNA immediately before electroporation. Cells were electroporated using the Neon system with three pulses at 1,500 V. After electroporation, cells were immediately transferred into pre-warmed X-VIVO 15 medium supplemented with 300 U ml⁻¹ IL-2, recovered at 37 °C for 30 min, and plated in 48-welL plates at 0.5–1 × 10⁶ cells ml⁻¹.

### Amplicon sequencing for CRISPR knock-in efficacy assay

Genomic DNA was extracted using the Quick DNA Miniprep Kit (Zymo) for each group from edited cells 3 days after electroporation. Genomic regions spanning the Cas9 cut sites were amplified using locus-specific primers **(Supplementary table 7)**, purified, indexed with Illumina-compatible adapters, pooled, and sequenced on a miniseq (Illumina). Amplicon sequencing reads were analyzed using CRISPResso2. Reads containing the expected donor-derived sequence were classified as HDR products, whereas reads containing indels at the Cas9 cut site without donor incorporation were classified as error-prone repair products. HDR editing efficiency was calculated as the percentage of aligned reads containing the expected donor-derived edit, and the HDR/error-prone repair ratio was calculated by dividing HDR reads by indel-containing non-HDR reads.

### Single cell PAIR-seq

HEK293T-PAIR KI cells stably expressing individual PAIR RNA constructs were generated by lentiviral transduction. 10 individual lenti-PAIR RNA constructs (P21, P53, P54, P56, P57, P61–P65, Supplementary Table 1) were separately transduced into HEK293T-PAIR cells. 48hours after infection, puromycin (10 µg/mL) was added for 24 hours to select transduced cells. Untraduced control cells showed no survival under selection. Following drug selection, the 10 individual PAIR-expressing cell populations were combined at comparable cell densities to generate a pooled PAIR cell population. The pooled cells were divided into six wells corresponding to two experimental groups, with three biological replicates per group. Group 1 cells were transfected with Cas9 ribonucleoprotein (RNP) complexed with sgRNA targeting the GAPDH locus (Cas9/sgGAPDH) using CRISPRMAX reagent. Group 2 cells were transfected with Cas9 RNP complexed with a non-targeting sgRNA (Cas9/sgNTC). For each biological replicate, 3125 ng of Cas9 protein and 625 ng of in vitro transcribed (IVT) sgRNA were delivered using CRISPRMAX reagent according to the manufacturer’s protocol (Invitrogen). 18 hours after transfection, approximately 1–2 × 10⁶ cells from each biological replicate were harvested and labeled individually using TotalSeq™ anti-human hashtag antibodies (C255, C256, C257) to enable sample multiplexing. For each experimental condition, three biological replicates were pooled after hashtag labeling and processed using the 10x Genomics Chromium 5′ v3 platform with a target cell recovery of 30,000 each group. Group 1 samples wereprocessed in 10x run #1, and Group 2 samples were processed in 10x run #2. For each 10x run, three libraries were generated: gene expression (mRNA), hashtag oligonucleotide (HTO), and PAIR RNA capture libraries. The libraries were then pooled at an appropriate ratio and sequenced on an Illumina NovaSeq X Platform.

### Single-cell RNA analysis

Cellular mRNA and HTO libraries were processed with Cell Ranger multi. PAIR direct-capture reads were processed using a custom four-step Python pipeline (code/PAIR_extraction/), including extraction of PAIR payload sequences from direct-capture reads, matching of cell barcodes to the mRNA whitelist, matching of PAIR barcodes to the PAIR-construct whitelist, and generation of a sparse PAIR UMI count matrix (cells × PAIR constructs). Ensembl gene identifiers in the mRNA count matrix were converted to HGNC symbols using biomaRt, and counts for duplicated gene symbols were summed. For PAIR tag assignment, each cell was assigned to the PAIR construct with the highest detected PAIR UMI count. Cells with zero detected PAIR UMIs were labeled as unassigned. For quality assessment, per-cell assignment metrics, including the top PAIR tag, top-tag UMI count, second-ranked tag, second-ranked UMI count, and assignment margin, were retained. Downstream analysis was performed in R using Seurat v5. For each sample, cells were filtered based on nCount_RNA, nFeature_RNA, and percent.mt, retaining cells within ±3 median absolute deviations of the sample-specific median. Expression values were log-normalized, and the top 2,000 highly variable genes were identified using the vst method. Data were scaled, principal component analysis was performed, and the top 20 principal components were used for shared nearest-neighbor graph construction, clustering, and UMAP visualization. Clustering was performed at a resolution of 0.4.

Differential expression and pathway analysis: For analysis of the NBN-conditioned DSB response (Fig. 4d), differential expression between RNP-treated and control cells was computed separately within the NT_CRISPRa and NBN_CRISPRa groups using Seurat FindMarkers with the Wilcoxon rank-sum test. For each gene, the NBN-conditioned DSB response score was defined as the difference between the corresponding log2 fold changes in the NBN_CRISPRa and NT_CRISPRa backgrounds. Genes were ranked by this score and analyzed by preranked gene set enrichment analysis using fgsea against the

MSigDB Hallmark collection, with Benjamini–Hochberg correction for multiple testing. For Tier 3 partner comparisons, differential expression between dual-perturbation cells (NBN + partner knockdown) and the NBN-only reference was performed within the RNP condition using Seurat FindMarkers with the Wilcoxon rank-sum test. Genes were ranked by differential-expression statistics and analyzed by pre-ranked fgsea against the Hallmark collection. Repair-pathway module scoring: Per-cell activity scores for curated homologous recombination (HR), non-homologous end joining (NHEJ), and microhomology-mediated end joining (MMEJ) gene sets were calculated using AUCell. Gene rankings were generated with AUCell_buildRankings, and enrichment scores were computed with AUCell_calcAUC using the default aucMaxRanksetting. AUC scores were z-scored across conditions for visualization. Residual and directional analyses: For analysis of the Module A RNP-context residual, a per-gene residual score was defined as FC_AB − FC_A, where FC_AB denotes the fold change of the dual perturbation relative to the wild-type reference under the RNP condition, and FC_A denotes the fold change of the NBN-only perturbation relative to the same reference. Genes with |residual| < 0.3 were classified as near-additive. Genes with |residual| ≥ 0.3 were classified according to the sign of the residual for descriptive stratification. For the directional opposition metric shown in Fig. 4h, genes were first selected from the NBN-conditioned DSB response set using an absolute score threshold of 0.25. For each partner knockdown, the fraction was calculated as the proportion of genes for which the fold change in the partner comparison within RNP-treated, NBN-activated cells (NBN + partner versus NBN + NT) had the opposite sign to the corresponding NBN-conditioned DSB response score. Cross-partner overlap analysis: Differentially expressed gene sets from Tier 3 partner comparisons were intersected across TP53BP1, XRCC6, and POLQ perturbations. Genes passing an adjusted P < 0.05 and |log2FC| > 0.5 threshold were used as the primary sets. When fewer than 20 genes passed this cutoff for a given comparison, a secondary threshold of nominal P < 0.05 and |log2FC| > 0.3 was used for overlap visualization. Set intersections were visualized with UpSetR, and pairwise similarity was quantified using the Jaccard index. All clustering, differential-expression, and enrichment analyses were performed on pooled single-cell data.

## References

1. Valls, P.O. & Esposito, A. Signalling dynamics, cell decisions, and homeostatic control in health and disease. Current Opinion in Cell Biology 75, 102066 (2022).

2. Liston, A. & Gray, D.H. Homeostatic control of regulatory T cell diversity. Nat Rev Immunol 14, 154–165 (2014).

3. Harapas, C.R. et al. Organellar homeostasis and innate immune sensing. Nat Rev Immunol 22, 535–549 (2022).

4. Herzog, H. Integrated pathways that control stress and energy homeostasis. Nat Rev Endocrinol 16, 75–76 (2020).

5. Chiolo, I., Altmeyer, M., Legube, G. & Mekhail, K. Nuclear and genome dynamics underlying DNA double-strand break repair. Nat Rev Mol Cell Biol (2025).

6. Wang, H., Yang, Y., Liu, J. & Qian, L. Direct cell reprogramming: approaches, mechanisms and progress. Nat Rev Mol Cell Biol 22, 410–424 (2021).

7. Nambiar, T.S., Baudrier, L., Billon, P. & Ciccia, A. CRISPR-based genome editing through the lens of DNA repair. Mol Cell 82, 348–388 (2022).

8. Southard, K.M. et al. Comprehensive transcription factor perturbations recapitulate fibroblast transcriptional states. Nat Genet 57, 2323–2334 (2025).

9. Magnusson, J.P. et al. PreciCE: Precision engineering of cell fates via data-driven multi-gene control of transcriptional networks. bioRxiv, 2024.2011.2004.621938 (2024).

10. Liu, C. et al. CRISPRi/a screens in human iPSC-cardiomyocytes identify glycolytic activation as a druggable target for doxorubicin-induced cardiotoxicity. Cell Stem Cell 31, 1760–1776 e1769 (2024).

11. Schmidt, R. et al. CRISPR activation and interference screens decode stimulation responses in primary human T cells. Science 375, eabj4008 (2022).

12. Knudsen, N.H. et al. In vivo CRISPR screens identify modifiers of CAR T cell function in myeloma. Nature 646, 953–962 (2025).

13. Mulvey, A., Trueb, L., Coukos, G. & Arber, C. Novel strategies to manage CAR-T cell toxicity. Nat Rev Drug Discov 24, 379–397 (2025).

14. Scully, R., Panday, A., Elango, R. & Willis, N.A. DNA double-strand break repair-pathway choice in somatic mammalian cells. Nat Rev Mol Cell Biol 20, 698–714 (2019).

15. Roidos, P. et al. A scalable CRISPR/Cas9-based fluorescent reporter assay to study DNA double-strand break repair choice. Nat Commun 11, 4077 (2020).

16. Gilbert, Luke A. et al. Genome-Scale CRISPR-Mediated Control of Gene Repression and Activation. Cell 159, 647–661 (2014).

17. Herken, B.W. et al. Large-scale mapping of environmental-genetic interactions illustrates the dynamic nature of cell-cycle and DNA repair regulation. Molecular Cell 86, 757–773.e755 (2026).

18. Burrell, W.H. et al. Rational design of synthetic proteins using a genome-scale CRISPR screen. bioRxiv, 2026.2002.2019.706875 (2026).

19. Fielden, J. et al. Comprehensive interrogation of synthetic lethality in the DNA damage response. Nature 640, 1093–1102 (2025).

20. Awwad, S.W., Serrano-Benitez, A., Thomas, J.C., Gupta, V. & Jackson, S.P. Revolutionizing DNA repair research and cancer therapy with CRISPR–Cas screens. Nature Reviews Molecular Cell Biology 24, 477–494 (2023).

21. Hussmann, J.A. et al. Mapping the genetic landscape of DNA double-strand break repair. Cell 184, 5653–5669 e5625 (2021).

22. Bowden, A.R. et al. Parallel CRISPR-Cas9 screens clarify impacts of p53 on screen performance. Elife 9 (2020).

23. Huang, M. et al. FACS-based genome-wide CRISPR screens define key regulators of DNA damage signaling pathways. Molecular Cell 83, 2810–2828.e2816 (2023).

24. Hartman, A. et al. A unified genetic perturbation language for human cellular programming. bioRxiv, 2025.2011.2020.689421 (2025).

25. Pacalin, N.M. et al. Bidirectional epigenetic editing reveals hierarchies in gene regulation. Nature Biotechnology (2024).

26. McCutcheon, S.R. et al. Transcriptional and epigenetic regulators of human CD8+ T cell function identified through orthogonal CRISPR screens. Nature Genetics 55, 2211–2223 (2023).

27. Shaw, W.M. et al. Inducible expression of large gRNA arrays for multiplexed CRISPRai applications. Nature Communications 13, 4984 (2022).

28. Drager, N.M. et al. A CRISPRi/a platform in human iPSC-derived microglia uncovers regulators of disease states. Nat Neurosci 25, 1149–1162 (2022).

29. Biering, S.B. et al. Genome-wide bidirectional CRISPR screens identify mucins as host factors modulating SARS-CoV-2 infection. Nature Genetics 54, 1078–1089 (2022).

30. Martella, A. et al. Systematic Evaluation of CRISPRa and CRISPRi Modalities Enables Development of a Multiplexed, Orthogonal Gene Activation and Repression System. ACS Synthetic Biology 8, 1998–2006 (2019).

31. Boettcher, M. et al. Dual gene activation and knockout screen reveals directional dependencies in genetic networks. Nature Biotechnology 36, 170–178 (2018).

32. Dahlman, J.E. et al. Orthogonal gene knockout and activation with a catalytically active Cas9 nuclease. Nature Biotechnology 33, 1159–1161 (2015).

33. Gao, Y. et al. Complex transcriptional modulation with orthogonal and inducible dCas9 regulators. Nat Methods 13, 1043–1049 (2016).

34. Truong, V.A. et al. CRISPRai for simultaneous gene activation and inhibition to promote stem cell chondrogenesis and calvarial bone regeneration. Nucleic Acids Research 47, e74–e74 (2019).

35. Replogle, J.M. et al. Mapping information-rich genotype-phenotype landscapes with genome-scale Perturb-seq. Cell 185, 2559–2575.e2528 (2022).

36. Norman, T.M. et al. Exploring genetic interaction manifolds constructed from rich single-cell phenotypes. Science 365, 786–793 (2019).

37. Konermann, S. et al. Genome-scale transcriptional activation by an engineered CRISPR-Cas9 complex. Nature 517, 583–588 (2015).

38. Pacesa, M., Pelea, O. & Jinek, M. Past, present, and future of CRISPR genome editing technologies. Cell 187, 1076–1100 (2024).

39. Konermann, S. et al. Transcriptome Engineering with RNA-Targeting Type VI-D CRISPR Effectors. Cell 173, 665–676 e614 (2018).

40. Zhang, C. et al. Structural Basis for the RNA-Guided Ribonuclease Activity of CRISPR-Cas13d. Cell 175, 212–223.e217 (2018).

41. Ali, S.E., Mittal, A. & Mathews, D.H. RNA Secondary Structure Analysis Using RNAstructure. Current Protocols 3, e846 (2023).

42. Chavez, A. et al. Highly efficient Cas9-mediated transcriptional programming. Nat Methods 12, 326–328 (2015).

43. Zhang, B. et al. Two HEPN domains dictate CRISPR RNA maturation and target cleavage in Cas13d. Nature Communications 10, 2544 (2019).

44. Nishimasu, H. et al. Crystal Structure of Cas9 in Complex with Guide RNA and Target DNA. Cell 156, 935–949 (2014).

45. East-Seletsky, A. et al. Two distinct RNase activities of CRISPR-C2c2 enable guide-RNA processing and RNA detection. Nature 538, 270–273 (2016).

46. Liu, L. et al. Two Distant Catalytic Sites Are Responsible for C2c2 RNase Activities. Cell 168, 121–134 e112 (2017).

47. Chang, C., Ma, G., Cheung, E. & Hutchins, A.P. A programmable system to methylate and demethylate N6-Methyladenosine (m6A) on specific RNA transcripts in mammalian cells. Journal of Biological Chemistry (2022).

48. Abramson, J. et al. Accurate structure prediction of biomolecular interactions with AlphaFold 3. Nature 630, 493–500 (2024).

49. Yang, Z.X. et al. Superior Fidelity and Distinct Editing Outcomes of SaCas9 Compared with SpCas9 in Genome Editing. Genomics Proteomics Bioinformatics 21, 1206–1220 (2023).

50. McCutcheon, S.R., Rohm, D., Iglesias, N. & Gersbach, C.A. Epigenome editing technologies for discovery and medicine. Nat Biotechnol 42, 1199–1217 (2024).

51. Kim, H.K. et al. In vivo high-throughput profiling of CRISPR-Cpf1 activity. Nat Methods 14, 153–159 (2017).

52. Domitrovich, A.M. & Kunkel, G.R. Multiple, dispersed human U6 small nuclear RNA genes with varied transcriptional efficiencies. Nucleic Acids Research 31, 2344–2352 (2003).

53. Chen, E. et al. Decorating chromatin for enhanced genome editing using CRISPR-Cas9. Proc Natl Acad Sci U S A 119, e2204259119 (2022).

54. Leuzzi, G. et al. SMARCAL1 is a dual regulator of innate immune signaling and PD-L1 expression that promotes tumor immune evasion. Cell (2024).

55. Deshpande, R.A. et al. Genome-wide analysis of DNA-PK-bound MRN cleavage products supports a sequential model of DSB repair pathway choice. Nat Commun 14, 5759 (2023).

56. Reint, G. et al. Rapid genome editing by CRISPR-Cas9-POLD3 fusion. Elife 10 (2021).

57. Su, D. et al. CRISPR/CAS9-based DNA damage response screens reveal gene-drug interactions. DNA Repair 87, 102803 (2020).

58. Chen, W. et al. Massively parallel profiling and predictive modeling of the outcomes of CRISPR/Cas9-mediated double-strand break repair. Nucleic Acids Res 47, 7989–8003 (2019).

59. Simpson, D., Ling, J., Jing, Y. & Adamson, B. Mapping the Genetic Interaction Network of PARP inhibitor Response. bioRxiv, 2023.2008.2019.553986 (2023).

60. Koblan, L.W., et al. Efficient C•G-to-G•C base editors developed using CRISPRi screens, target-library analysis, and machine learning. Nature Biotechnology 39, 1414–1425 (2021).

61. Chen, P.J. et al. Enhanced prime editing systems by manipulating cellular determinants of editing outcomes. Cell 184, 5635–5652.e5629 (2021).

62. Adamson, B., Smogorzewska, A., Sigoillot, F.D., King, R.W. & Elledge, S.J. A genome-wide homologous recombination screen identifies the RNA-binding protein RBMX as a component of the DNA-damage response. Nat Cell Biol 14, 318–328 (2012).

63. Cullot, G. et al. Genome editing with the HDR-enhancing DNA-PKcs inhibitor AZD7648 causes large-scale genomic alterations. Nat Biotechnol (2024).

64. Rees, H.A., Yeh, W.H. & Liu, D.R. Development of hRad51-Cas9 nickase fusions that mediate HDR without double-stranded breaks. Nat Commun 10, 2212 (2019).

65. Carusillo, A. et al. A novel Cas9 fusion protein promotes targeted genome editing with reduced mutational burden in primary human cells. Nucleic Acids Res 51, 4660–4673 (2023).

66. Charpentier, M. et al. CtIP fusion to Cas9 enhances transgene integration by homology-dependent repair. Nat Commun 9, 1133 (2018).

67. Canny, M.D. et al. Inhibition of 53BP1 favors homology-dependent DNA repair and increases CRISPR-Cas9 genome-editing efficiency. Nat Biotechnol 36, 95–102 (2018).

68. Arnoult, N. et al. Regulation of DNA repair pathway choice in S and G2 phases by the NHEJ inhibitor CYREN. Nature 549, 548–552 (2017).

69. Jayavaradhan, R. et al. CRISPR-Cas9 fusion to dominant-negative 53BP1 enhances HDR and inhibits NHEJ specifically at Cas9 target sites. Nat Commun 10, 2866 (2019).

70. Tran, N.T. et al. Enhancement of Precise Gene Editing by the Association of Cas9 With Homologous Recombination Factors. Front Genet 10, 365 (2019).

71. Paulsen, B.S. et al. Ectopic expression of RAD52 and dn53BP1 improves homology-directed repair during CRISPR-Cas9 genome editing. Nat Biomed Eng 1, 878–888 (2017).

72. Savic, N. et al. Covalent linkage of the DNA repair template to the CRISPR-Cas9 nuclease enhances homology-directed repair. Elife 7 (2018).

73. Pinder, J., Salsman, J. & Dellaire, G. Nuclear domain’knock-in’ screen for the evaluation and identification of small molecule enhancers of CRISPR-based genome editing. Nucleic Acids Res 43, 9379–9392 (2015).

74. Wang, J.Y. & Doudna, J.A. CRISPR technology: A decade of genome editing is only the beginning. Science 379, eadd8643 (2023).

75. Gallagher, D.N. & Haber, J.E. Repair of a Site-Specific DNA Cleavage: Old-School Lessons for Cas9-Mediated Gene Editing. Acs Chem Biol 13, 397–405 (2018).

76. Falbo, L. & Costanzo, V. Replicative gaps in DNA damage tolerance, genome instability, and cancer therapy. Mol Cell (2026).

77. Bantele, S. et al. Repair of DNA double-strand breaks leaves heritable impairment to genome function. Science 390, eadk6662 (2025).

78. van de Kooij, B., Kruswick, A., van Attikum, H. & Yaffe, M.B. Multi-pathway DNA-repair reporters reveal competition between end-joining, single-strand annealing and homologous recombination at Cas9-induced DNA double-strand breaks. Nat Commun 13, 5295 (2022).

79. Chen, S. et al. Structural basis of long-range to short-range synaptic transition in NHEJ. Nature 593, 294–298 (2021).

80. Sun, Y.R., McCorvie, T.J., Yates, L.A. & Zhang, X.D. Structural basis of homologous recombination. Cellular and Molecular Life Sciences 77, 3–18 (2020).

81. Symington, L.S. & Gautier, J. Double-strand break end resection and repair pathway choice. Annu Rev Genet 45, 247–271 (2011).

82. Gilbert, L.A. et al. Genome-Scale CRISPR-Mediated Control of Gene Repression and Activation. Cell 159, 647–661 (2014).

83. Blackford, A.N. & Jackson, S.P. ATM, ATR, and DNA-PK: The Trinity at the Heart of the DNA Damage Response. Mol Cell 66, 801–817 (2017).

84. Ciccia, A. & Elledge, S.J. The DNA damage response: making it safe to play with knives. Mol Cell 40, 179–204 (2010).

85. Glaser, A., McColl, B. & Vadolas, J. GFP to BFP Conversion: A Versatile Assay for the Quantification of CRISPR/Cas9-mediated Genome Editing. Molecular Therapy - Nucleic Acids 5, e334 (2016).

86. Rotheneder, M. et al. Cryo-EM structure of the Mre11-Rad50-Nbs1 complex reveals the molecular mechanism of scaffolding functions. Mol Cell 83, 167–185 e169 (2023).

87. Deshpande, R.A., Lee, J.H., Arora, S. & Paull, T.T. Nbs1 Converts the Human Mre11/Rad50 Nuclease Complex into an Endo/Exonuclease Machine Specific for Protein-DNA Adducts. Mol Cell 64, 593–606 (2016).

88. You, Z. et al. CtIP Links DNA Double-Strand Break Sensing to Resection. Molecular Cell 36, 954–969 (2009).

89. Zhao, F., Kim, W., Kloeber, J.A. & Lou, Z. DNA end resection and its role in DNA replication and DSB repair choice in mammalian cells. Exp Mol Med 52, 1705–1714 (2020).

90. Kim, H.K. et al. In vivo high-throughput profiling of CRISPR–Cpf1 activity. Nature Methods 14, 153–159 (2017).

91. Clement, K. et al. CRISPResso2 provides accurate and rapid genome editing sequence analysis. Nature Biotechnology 37, 224–226 (2019).

92. Datlinger, P. et al. Pooled CRISPR screening with single-cell transcriptome readout. Nature Methods 14, 297–301 (2017).

93. Dixit, A. et al. Perturb-Seq: Dissecting Molecular Circuits with Scalable Single-Cell RNA Profiling of Pooled Genetic Screens. Cell 167, 1853–1866.e1817 (2016).

94. Adamson, B. et al. A Multiplexed Single-Cell CRISPR Screening Platform Enables Systematic Dissection of the Unfolded Protein Response. Cell 167, 1867–1882 e1821 (2016).

95. Replogle, J.M. et al. Combinatorial single-cell CRISPR screens by direct guide RNA capture and targeted sequencing. Nat Biotechnol 38, 954–961 (2020).

96. Jin, X. et al. In vivo Perturb-Seq reveals neuronal and glial abnormalities associated with autism risk genes. Science 370, eaaz6063 (2020).

97. Lachmann, A., Xie, Z. & Ma’ayan, A. blitzGSEA: efficient computation of gene set enrichment analysis through gamma distribution approximation. Bioinformatics 38, 2356–2357 (2022).

98. Subramanian, A. et al. Gene set enrichment analysis: A knowledge-based approach for interpreting genome-wide expression profiles. Proceedings of the National Academy of Sciences 102, 15545–15550 (2005).

99. Liberzon, A. et al. The Molecular Signatures Database (MSigDB) hallmark gene set collection. Cell Syst 1, 417–425 (2015).

100. Panier, S. & Boulton, S.J. Double-strand break repair: 53BP1 comes into focus. Nat Rev Mol Cell Biol 15, 7–18 (2014).

101. Aibar, S. et al. SCENIC: single-cell regulatory network inference and clustering. Nat Methods 14, 1083–1086 (2017).

102. Eyquem, J. et al. Targeting a CAR to the TRAC locus with CRISPR/Cas9 enhances tumour rejection. Nature 543, 113–117 (2017).

103. Roth, T.L. et al. Reprogramming human T cell function and specificity with non-viral genome targeting. Nature 559, 405–409 (2018).

104. Richardson, C.D., Ray, G.J., DeWitt, M.A., Curie, G.L. & Corn, J.E. Enhancing homology-directed genome editing by catalytically active and inactive CRISPR-Cas9 using asymmetric donor DNA. Nat Biotechnol 34, 339–344 (2016).

105. Zetsche, B. et al. Multiplex gene editing by CRISPR-Cpf1 using a single crRNA array. Nat Biotechnol 35, 31–34 (2017).

106. Villiger, L. et al. CRISPR technologies for genome, epigenome and transcriptome editing. Nature Reviews Molecular Cell Biology (2024).

107. Foy, S.P. et al. Non-viral precision T cell receptor replacement for personalized cell therapy. Nature 615, 687–696 (2023).

108. Xie, K. et al. Efficient non-viral immune cell engineering using circular single-stranded DNA-mediated genomic integration. Nature Biotechnology 43, 1821–1832 (2025).

109. Roth, T.L. et al. Non-viral intron knock-ins for targeted gene integration into human T cells and for T-cell selection. Nature Biomedical Engineering (2025).

110. Allen, A.G. et al. A highly efficient transgene knock-in technology in clinically relevant cell types. Nature Biotechnology 42, 458–469 (2024).

111. Shy, B.R. et al. High-yield genome engineering in primary cells using a hybrid ssDNA repair template and small-molecule cocktails. Nature Biotechnology 41, 521–531 (2023).

112. Nyberg, W.A. et al. In vivo site-specific engineering to reprogram T cells. Nature 652, 712–721 (2026).

113. An, J. et al. Enhancement of the viability of T cells electroporated with DNA via osmotic dampening of the DNA-sensing cGAS-STING pathway. Nat Biomed Eng 8, 149–164 (2024).

114. Crinier, A., Narni-Mancinelli, E., Ugolini, S. & Vivier, E. SnapShot: Natural Killer Cells. Cell 180, 1280–1280 e1281 (2020).

115. Tsuchida, C.A. et al. Mitigation of chromosome loss in clinical CRISPR-Cas9-engineered T cells. Cell 186, 4567–4582 e4520 (2023).

116. Stadtmauer, E.A. et al. CRISPR-engineered T cells in patients with refractory cancer. Science 367, eaba7365 (2020).

117. Ray, U., Vartak, S.V. & Raghavan, S.C. NHEJ inhibitor SCR7 and its different forms: Promising CRISPR tools for genome engineering. Gene 763, 144997 (2020).

118. Srivastava, M. et al. An Inhibitor of Nonhomologous End-Joining Abrogates Double-Strand Break Repair and Impedes Cancer Progression. Cell 151, 1474–1487 (2012).

119. Richardson, R.R. et al. Enhancing Precision and Efficiency of Cas9-Mediated Knockin Through Combinatorial Fusions of DNA Repair Proteins. The CRISPR Journal 6, 447–461 (2023).

120. Li, Z. et al. Intrinsic targeting of host RNA by Cas13 constrains its utility. Nature Biomedical Engineering 8, 177–192 (2024).

121. Hart, S.K. et al. Precise RNA targeting with CRISPR–Cas13d. Nature Biotechnology 44, 64–69 (2026).

