## Supplementary Figures for "Parallel Activation and Interference CRISPR (PAIR) with Sequencing Uncovers DNA Repair Networks Guiding Precision Cell Engineering"

### Extended Data Fig. 1 (Related to Fig. 1)

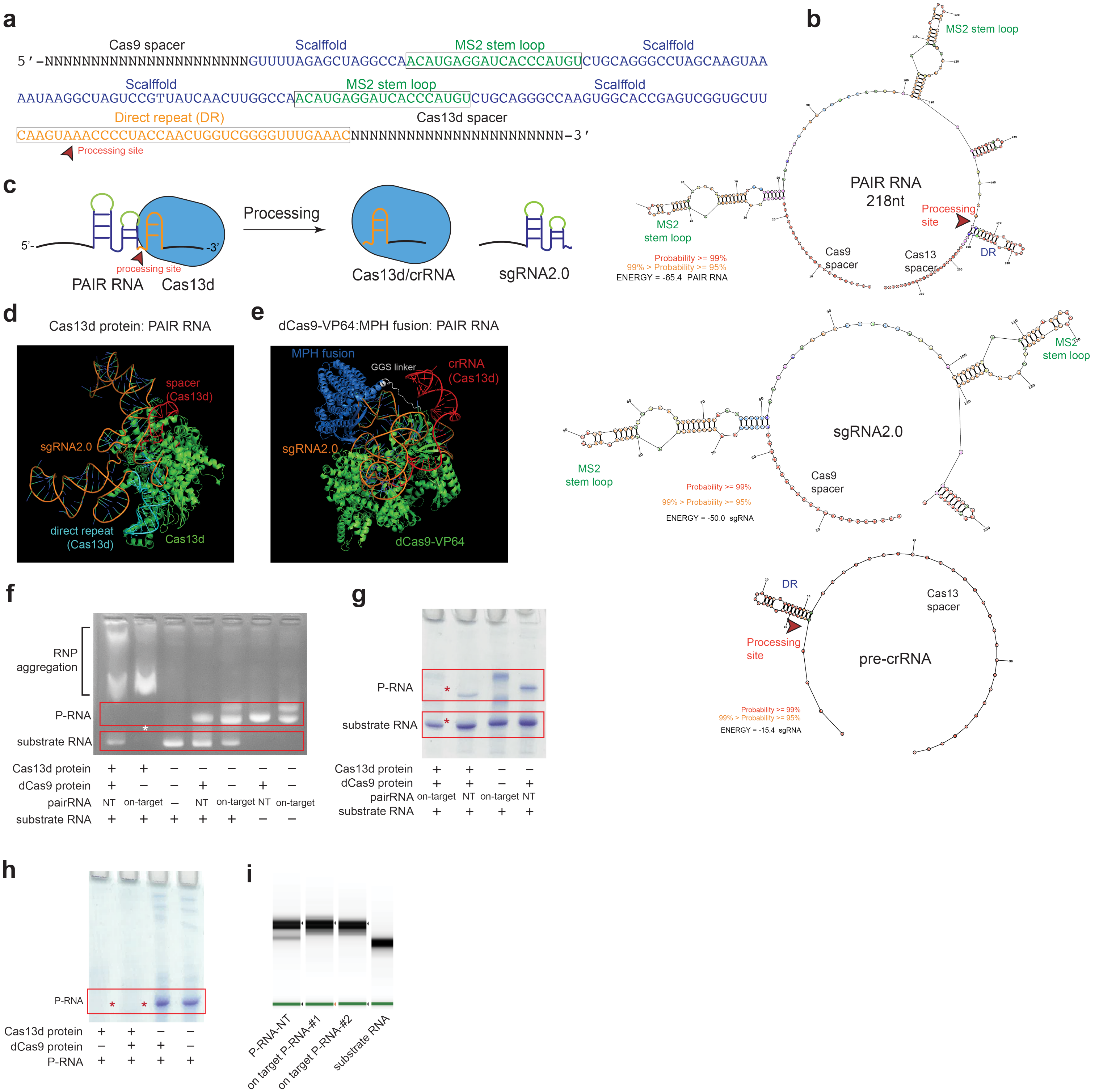


**Extended Data Fig.1 | (related to Fig.1). Establishing PAIR for bidirectional perturbation of two genes in single cells. a,** General sequence of the PAIR RNA. Distinct sequence elements are color-coded, including the spacer, scaffold, and MS2 stem loops in the sgRNA2.0 module, and the direct repeat (DR) and spacer in the pre-crRNA module. The predicted Cas13d processing site within the pre-crRNA region is indicated by a red triangle. **b,** Predicted secondary structures of the full-length PAIR RNA, sgRNA2.0, and pre-crRNA. The PAIR RNA prediction showed preservation of the expected local structural features of both guide modules, including the MS2 stem loops and direct repeat (DR). **c,** Schematic of Cas13d-mediated processing of the PAIR RNA. Cas13d (blue) recognizes the pre-crRNA DR region (orange) within the tandem PAIR RNA and processes it at the predicted processing site (red triangle). The processed crRNA then associates with Cas13d to guide target RNA interference, whereas the sgRNA2.0 module remains available for CRISPRa-mediated gene activation. **d,** AlphaFold3-predicted structural model of Cas13d bound to the PAIR RNA. The model shows Cas13d recognizing the 3′ pre-crRNA region of the PAIR RNA, including the Cas13d spacer and direct repeat, while the sgRNA2.0 portion remains positioned outside the Cas13d-bound region. **e,** AlphaFold3-predicted structural model of the CRISPRa module bound to the PAIR RNA. The model shows the 5′ sgRNA2.0 region associated with dCas9–VP64 and the MS2 stem-loop elements engaged by the MPH fusion protein, supporting recruitment of the CRISPRa effector complex through the PAIR RNA. **f,** Agarose gel analysis of in vitro Cas13d-mediated PIAR RNA (P-RNA) processing and substrate RNA digestion. IVT-produced P-RNA was incubated with Cas13d and/or dCas9 protein in the presence of on-target or non-target substrate RNA. Red boxes indicate P-RNA (up) and substrate RNA (bottom) species. **g,** Native PAGE analysis of Cas13d-mediated substrate RNA digesting with on-targeting P-RNA. On-target PAIR RNA supported substrate RNA cleavage in the presence of Cas13d, whereas non-target PAIR RNA or reactions lacking Cas13d did not produce comparable substrate cleavage. **h,** Native PAGE analysis of PAIR RNA processing by Cas13d in the presence or absence of dCas9 protein. Red boxes indicate P-RNA species, showing Cas13d-dependent processing of P-RNA under the indicated RNP assembly conditions. **i,** RNA Tapestation result of T7 IVT generated P-RNAs and substrate RNA.

### Extended Data Fig. 2 (Related to Fig. 1)

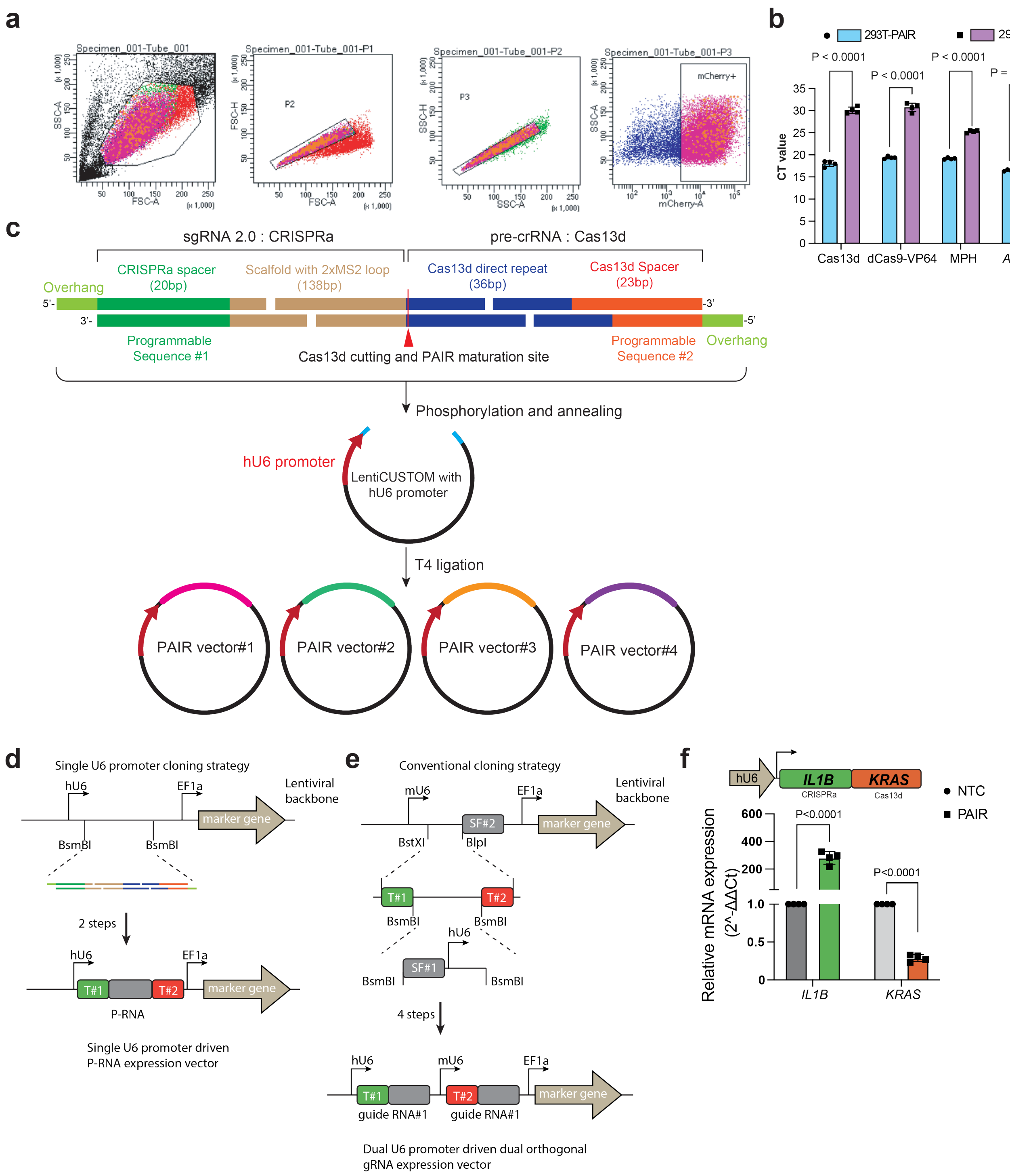


**Extended Data Fig.2 | (related to Fig.1). Establishing PAIR for bidirectional perturbation of two genes in single cells. a,** Flow-cytometry gating strategy used to enrich PAIR-positive HEK293T cells by FACS. **b,** RT–qPCR in 293T-PAIR cells showing Cas13d, dCas9, and MPH expression compared with wild-type HEK293T cells. Points indicate biological replicates; bars indicate mean (n = 4 biological replicates; two technical replicates each). **c,**Schematic of the cloning strategy for generating PAIR expression vectors. **d,** Schematics of single U6 promoter driven P-RNA vector workflow. **e,** Schematics of the conventional dual U6 promoter driven dual orthogonal guide RNA vector workflow. **f,** RT–qPCR showing simultaneous activation and interference by PAIR (IL1B up, KRAS down) in 293T-PAIR cells. Points indicate biological replicates; bars indicate mean (n = 4 biological replicates; two technical replicates each). P values were calculated by one-way ANOVA.

### Extended Data Fig. 3 (Related to Figs. 2)

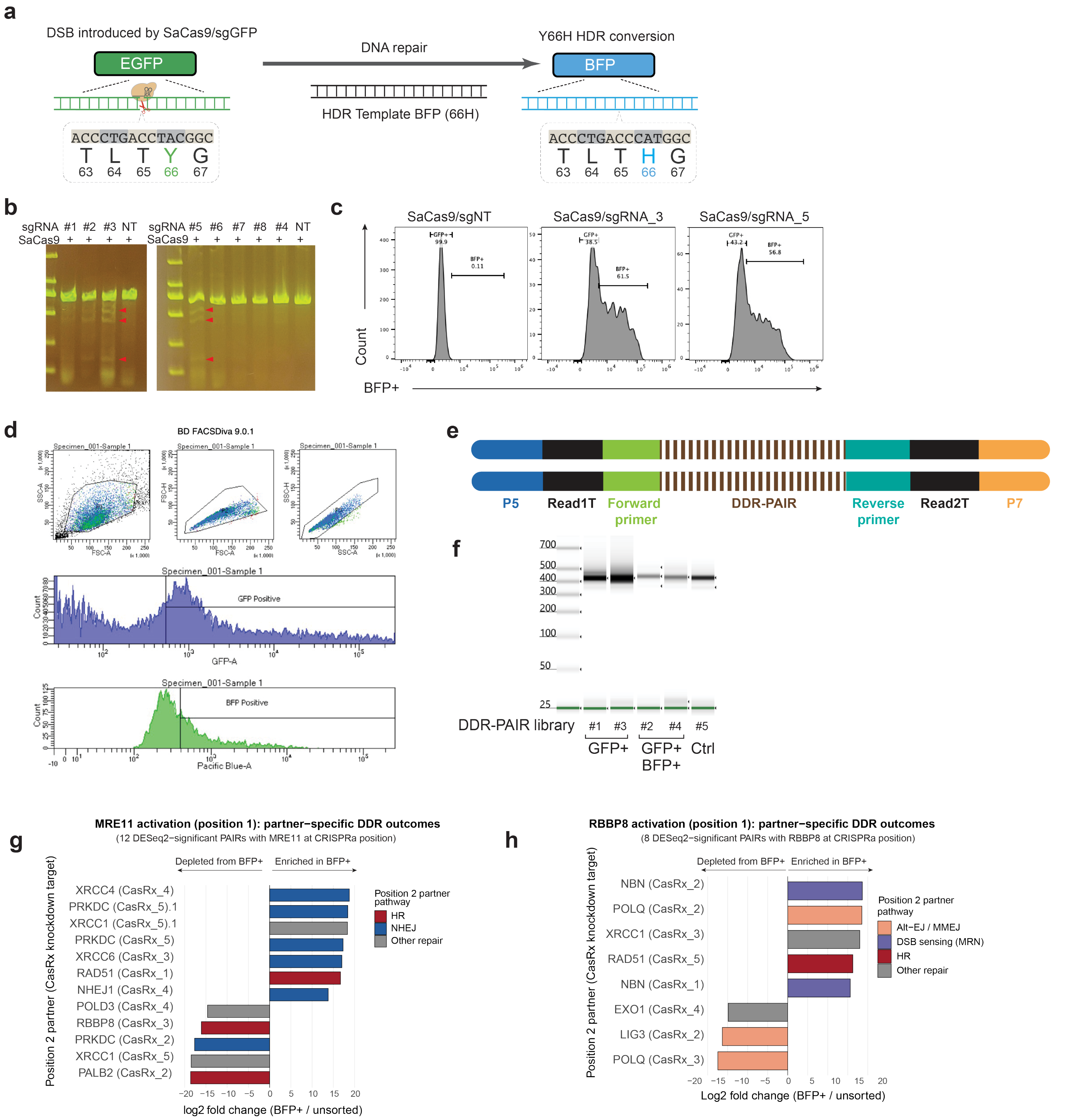


**Extended Data Fig. 3 | (related to Fig.2) PAIR-seq screen enables high-throughput dissection of competing DDR pathways**. **a,** Schematic of the GFP-to-BFP HDR reporter. SaCas9 introduces a DSB at the EGFP(66Y) locus and an HDR donor carrying the BFP(66H) sequence enables template-directed repair, yielding BFP signal. **b,** T7EN1 assay of cutting efficiency, Genomic PCR across the reporter locus following mRNA SaCas9/sgRNA delivery using the indicated sgRNAs (NT, non-targeting control). Red arrowheads indicate the expected amplicon/edited products. **c,** Representative flow-cytometry histograms of BFP signal in cells transfected with non-targeting sgRNA (sgNT) or the indicated sgRNAs together with the HDR donor.**d,** Gating strategy for FACS enrichment of reporter-positive populations. Representative plots and histograms show GFP positive and BFP positive gates used for downstream analysis. **e,** Schematics of DDR-PAIR-seq library structure. **f,** PCR amplification of integrated PAIR-DDR library cassettes from sorted populations (GFP+ and GFP+ BFP+) and control samples, confirming successful recovery of library sequences for NGS. The expected library size is 423bp. **g,** Representative independent validation of DDR-PAIR-seq hits by measuring target gene expressions using qRT-PCR. Dots represent biological replicates; bars indicate mean ± s.d. (n = 4 biological replicates, each with two technical replicates). P values were calculated using one-way ANOVA.

### Extended Data Fig. 4 (Related to Fig. 2, 3)

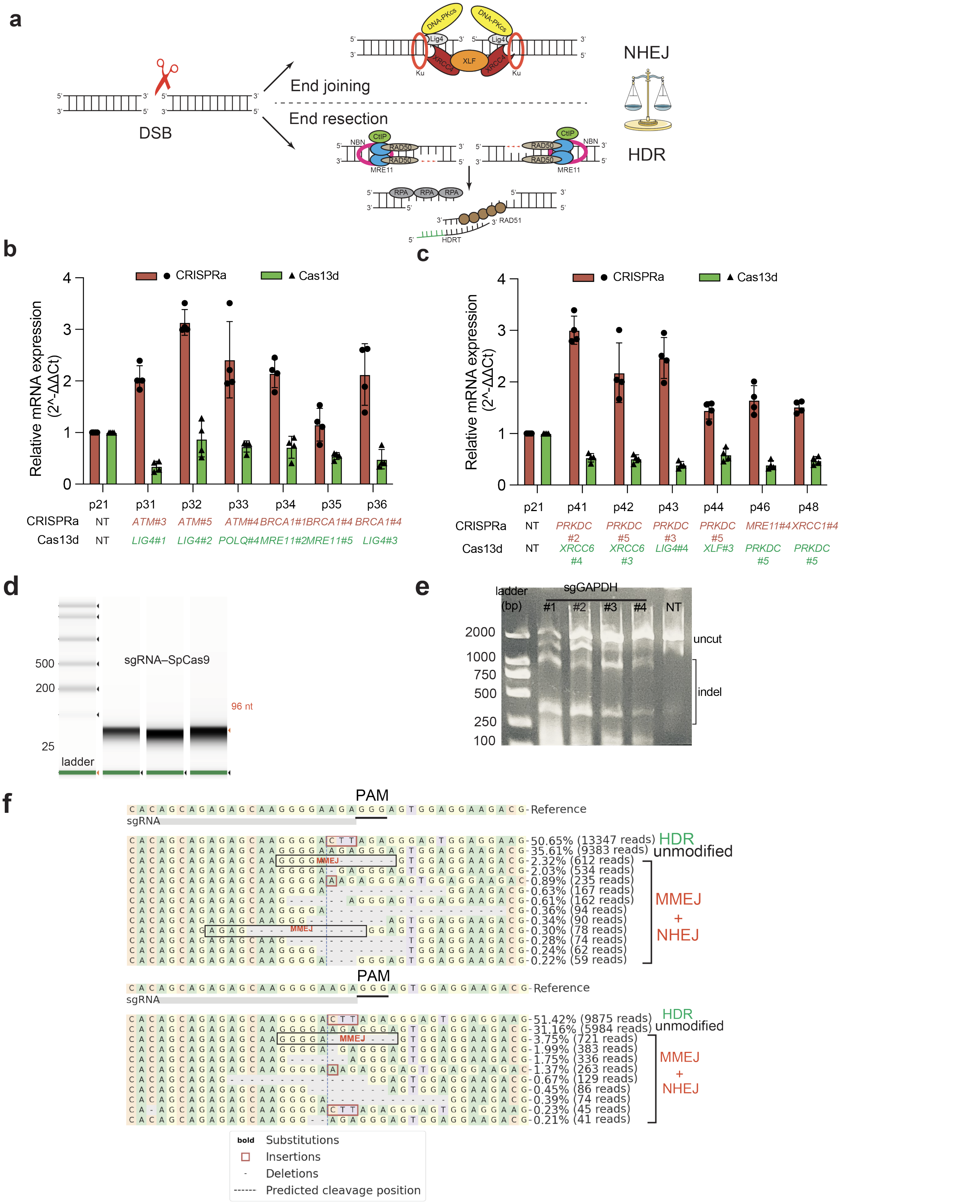


**Extended Data Fig.4 | (relative to Fig.2, 3) Validation of top PAIR combinations from DDR-seq**. **a,** Schematic of the two major DNA double-strand break (DSB) repair pathways. Following DSB induction, broken ends are channeled into one of two competing pathways. In non-homologous end joining (NHEJ; top), the Ku70/80 heterodimer rapidly binds DNA ends and recruits DNA-PKcs, followed by the XRCC4–XLF–LIG4 ligation complex, which directly re-ligates the two ends with little or no sequence homology, often introducing small insertions or deletions. In homology-directed repair (HDR; bottom), the MRE11–RAD50–NBN (MRN) complex together with CtIP initiates 5′ →3′ end resection to generate 3′ single-stranded DNA (ssDNA) overhangs. The resulting ssDNA is first coated by RPA and subsequently replaced by RAD51, forming a nucleoprotein filament that invades a homologous donor template (HDR template, HDRT) to enable templated, precise repair. **b, c,** Independent validation of DDR-PAIR-seq hits by measuring target gene expressions using RT-qPCR, Dots represent biological replicates, and bars represent the mean of n = 4 biological replicates, each with two technical replicates, p Value is from a one-way ANOVA. **d,** T7 IVT generated sgRNAs for RNP transfection. The expected sgRNA sizes are 96nt. **e,**T7EN1 assay for GAPDH sgRNA cutting efficiency test. **f,**T7EN1 assay for TRAC sgRNA cutting efficiency test. **f,** Representative CRISPResso2 alignment plots showing HDR and error-prone repair outcomes at the AAVS1 locus.

### Extended Data Fig. 5 (Related to Fig. 4)

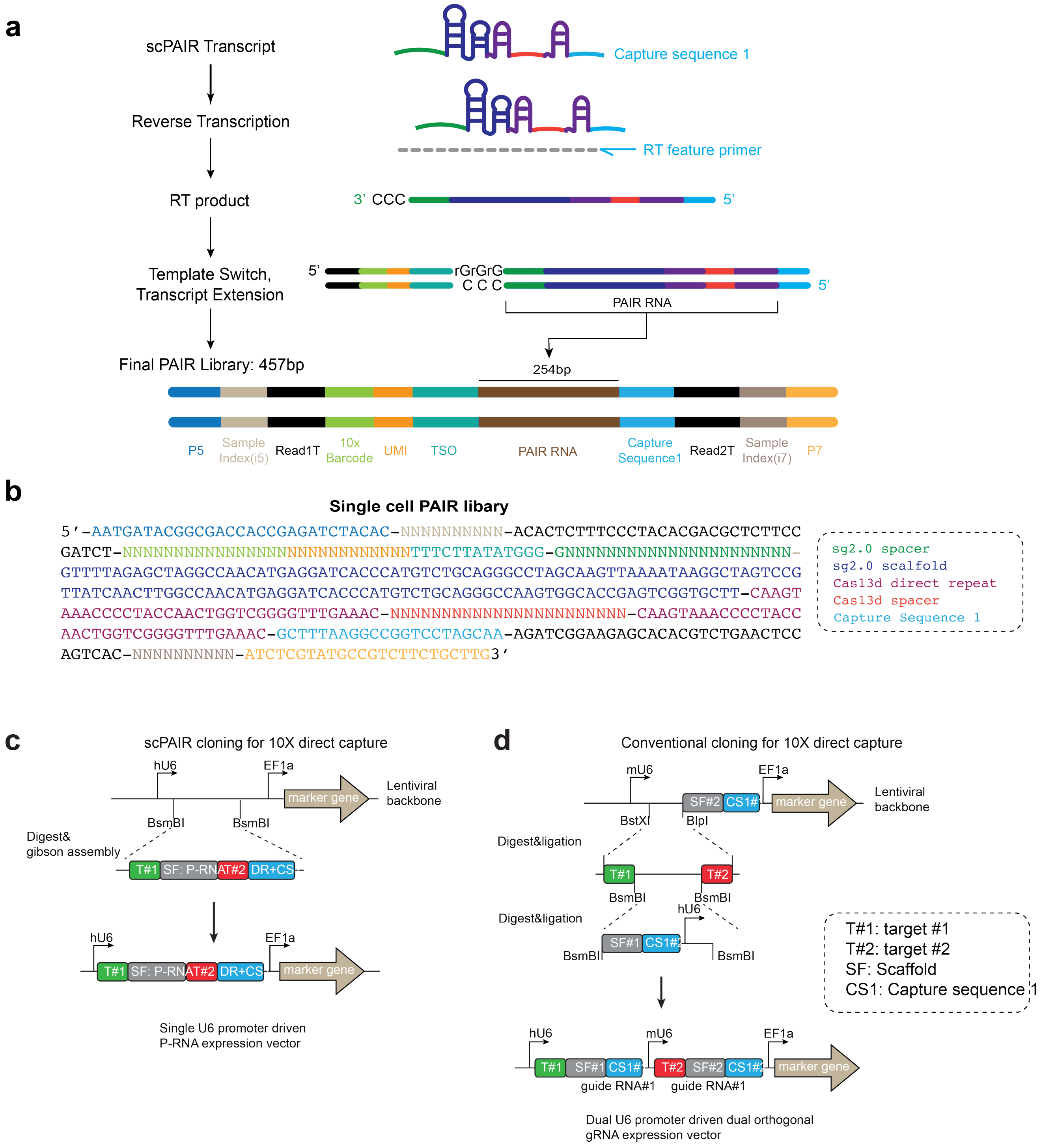


**Extended Data Fig. 5 | (related to Fig.4). Design of single cell PAIR-seq library. a,** Schematic of the scPAIR-seq transcript capture and reverse-transcription strategy. The 3′ capture sequence 1 (CS1) was appended downstream of the PAIR RNA to enable direct capture during 10x Genomics 5′ single-cell library construction. After reverse transcription with the feature RT primer and template switching, the final PAIR library contains the cell barcode, UMI, PAIR RNA sequence and capture handle for perturbation assignment. **b,** Sequence of single-cell PAIR RNA. **c,** Schematic of the single-U6-driven scPAIR vector design for direct PAIR RNA capture by 10x Genomics. **d,** Schematic of a conventional dual-U6 promoter strategy for 10x direct capture of dual orthogonal guide RNAs.

### Extended Data Fig. 6 (Related to Fig. 4)

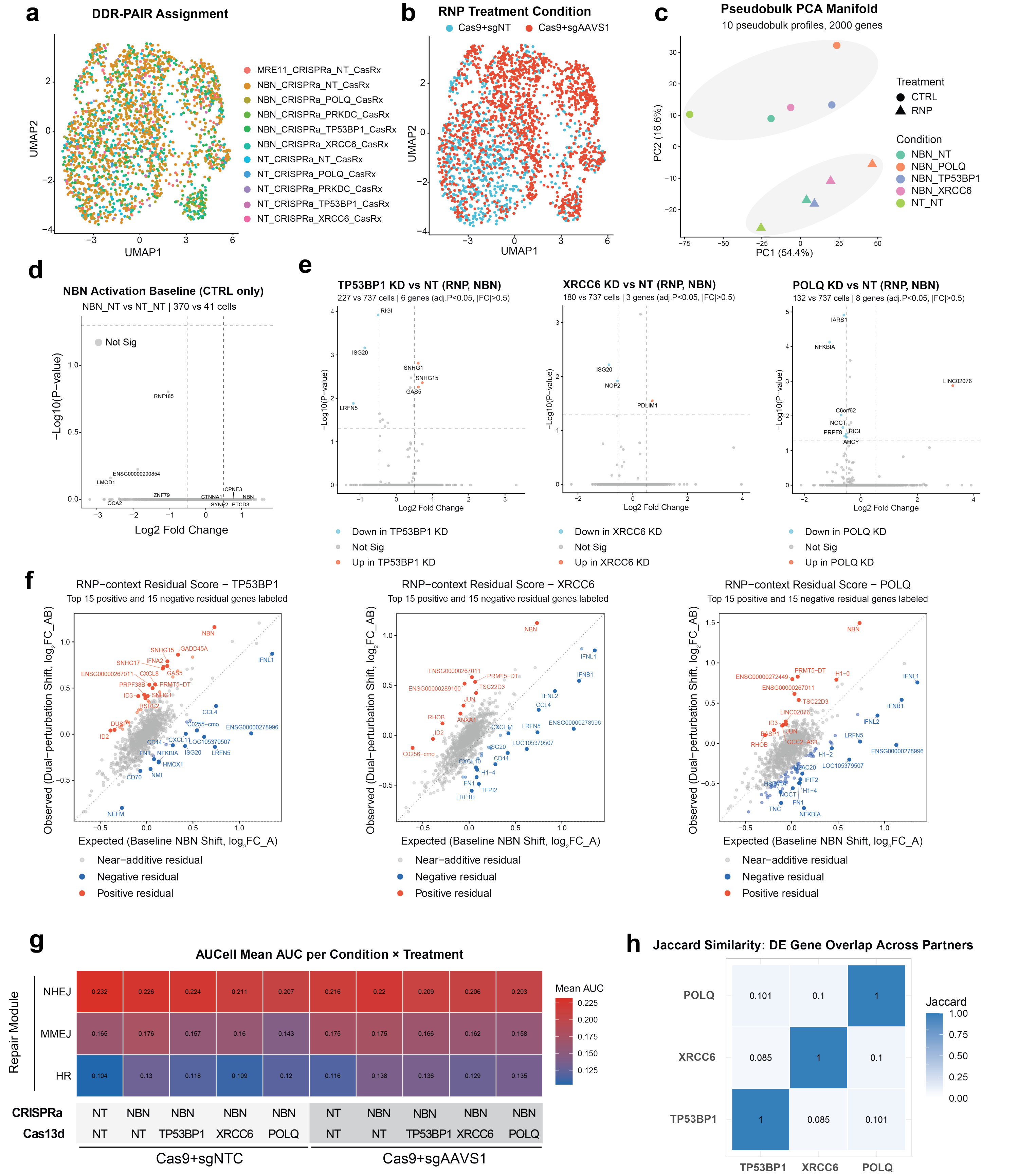

**Extended Data Fig. 6 | (related to Fig. 4) Single cell perturbation assignment, and supporting analyses for scPAIR-seq of the NBN interaction axis. a,** Uniform manifold approximation and projection (UMAP) embedding of the 2,271 post-QC cells colored by assigned PAIR identity. **b,** Same UMAP colored by RNP vs CTRL treatment condition. **c,** Pseudobulk principal-component analysis across perturbation arms and treatments; the first principal component separates RNP from CTRL cells across every arm, indicating that Cas9-induced DSB is the dominant axis of transcriptional variation. **d,** Volcano plot of the NBN_CRISPRa vs NT_CRISPRa contrast within the CTRL (no-RNP) condition. No genes pass adj. $P < 0.05$ and $|\log_2FC| > 0.5$ across 11,610 tested features. **e,** Volcano plots of each partner knockdown (TP53BP1, XRCC6, POLQ) versus the NT reference within the NBN-activated RNP condition. Cell-level Wilcoxon rank-sum; points colored by direction. **f,** RNP-context residual analysis (residual = $\log_2FC\sim AB - \log_2FC\sim A$) per partner. Most genes fall within ±0.3 of the NBN-only fold change (Near-additive residual; TP53BP1 97.6%, XRCC6 98.2%, POLQ 95.8% of 2,000 genes), with a tail of Positive- and Negative-residual genes, enabled by the bidirectional (activation + suppression) geometry of PAIR perturbation. **g,** AUCell mean AUC per condition × treatment (z-scored across conditions) for the HR, NHEJ, and MMEJ gene-set modules. **h,** Jaccard similarity of the partner-knockdown differentially expressed gene sets (cell-level Wilcoxon, adj. $P < 0.05$, $|\log_2FC| > 0.5$).

### Extended Data Fig. 7 (Related to Fig. 5)

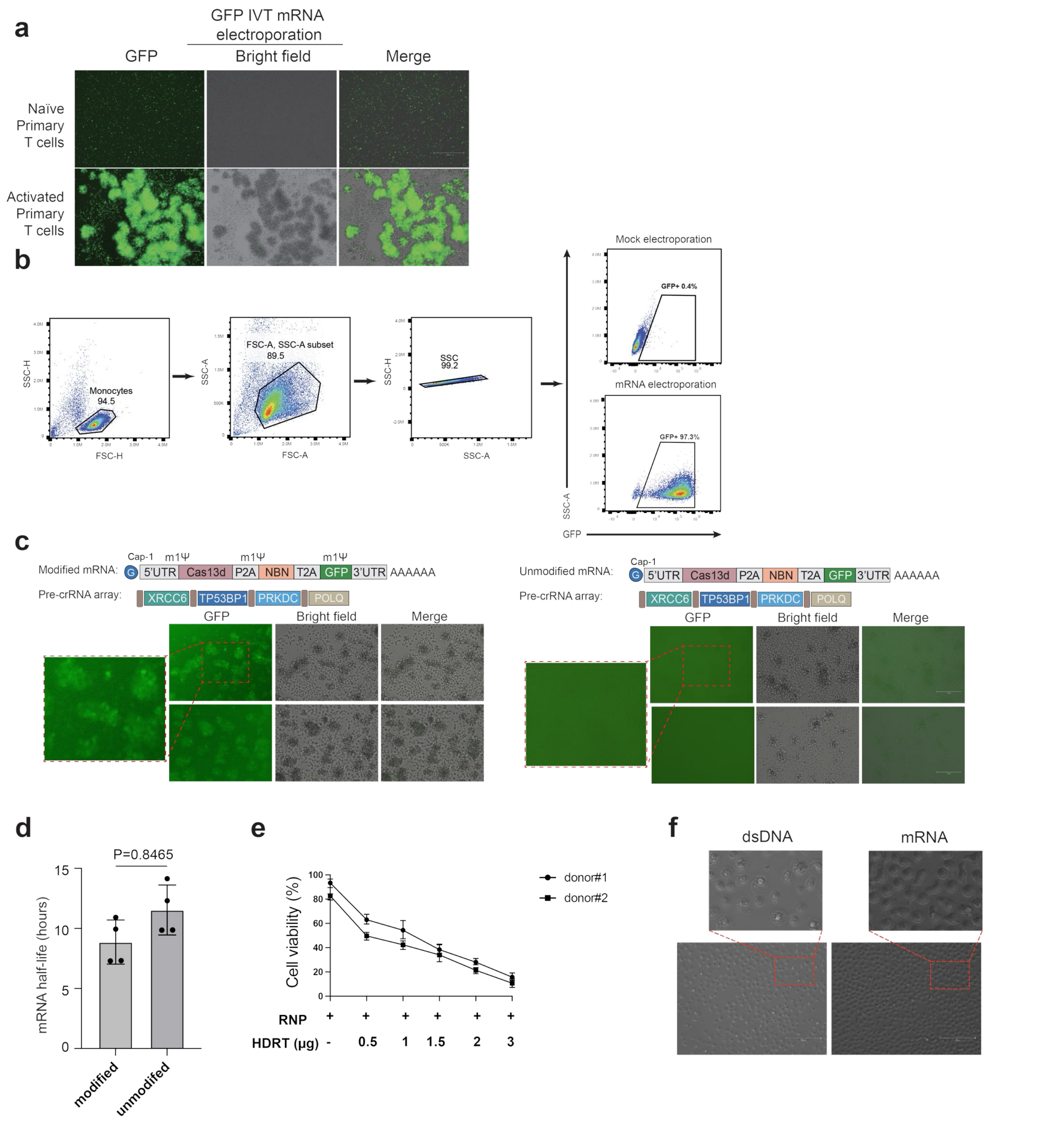


**Extended Data Fig. 7 | (raleted to Fig. 5) POKER-based bidirectional RNA regulation improves non-viral CAR-T engineering efficiency. a,** Quantification of mCherry-positive cells in the GAPDH knock-in assay with or without POKER priming. Each dot represents one independent electroporation replicate; bars indicate mean. Statistical significance was assessed as indicated (P values shown). **b,** Representative flow-cytometry gating strategy for analysis of electroporated primary T cells and validation of GFP mRNA delivery, including exclusion of debris and doublets and identification of GFP-positive cells. **c,** Representative fluorescence and bright-field images comparing capped/chemically modified POKER mRNA versus uncapped/unmodified mRNA (with the same pre-crRNA array). Robust GFP signal is observed with the capped/modified POKER mRNA, whereas little to no GFP fluorescence is detected with the uncapped/unmodified construct. **d,** Estimated mRNA half-life in primary T cells for capped/-modified versus uncapped/unmodified POKER mRNA constructs (mean ± s.e.m.; P value as indicated), each dots indicate individual donor. **e,** Cell viability of primary T cells following Cas9 RNP electroporation with increasing amounts of dsDNA HDR template (HDRT) across two human donors, illustrating a dose-dependent reduction in viability at higher donor DNA doses. **f,** Representative bright-field images comparing primary T cells after electroporation with dsDNA versus mRNA, illustrating reduced visible cellular stress and improved morphology following mRNA delivery relative to dsDNA under the conditions tested.

### Extended Data Fig. 8 (Related to Fig. 5)

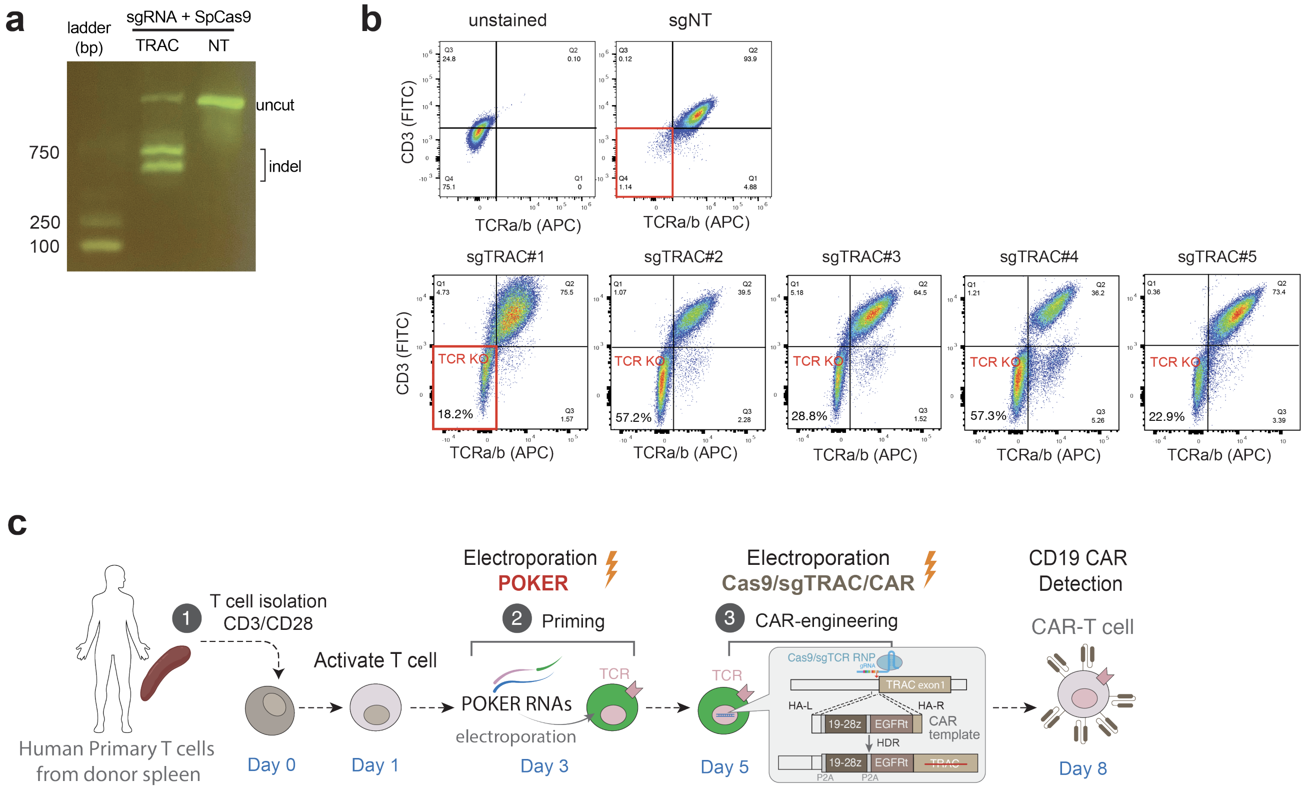


**Extended Data Fig. 8 | (raleted to Fig. 5) POKER-based bidirectional RNA regulation improves non-viral CAR-T engineering efficiency. a,** Agarose gel analysis of TRAC locus disruption following electroporation of primary human T cells with SpCas9 RNP and sgRNAs targeting TRAC (representative uncut and indel bands shown), compared with a non-targeting control. **b,** Flow-cytometry screen of candidate sgTRAC guides for efficient TCR disruption in primary human T cells. Representative plots show CD3 and TCRαβ staining following electroporation with the indicated sgRNAs, identifying guides that effectively reduce surface TCR expression for downstream knock-in engineering. **c,** Schematic of the POKER-primed workflow for non-viral TRAC CAR knock-in in primary human T cells, including T-cell isolation and activation, POKER RNA priming, subsequent Cas9/sgTRAC RNP plus dsDNA CAR HDR template electroporation, and downstream CAR detection.
